# Dynamic Functional Brain Reconfiguration During Sustained Pain

**DOI:** 10.1101/2021.10.16.464642

**Authors:** Jae-Joong Lee, Sungwoo Lee, Dong Hee Lee, Choong-Wan Woo

## Abstract

Pain is constructed through complex interactions among multiple brain systems, but it remains unclear how functional brain network representations are dynamically reconfigured over time while experiencing pain. Here, we investigated the dynamic changes in the functional brain networks during 20-min capsaicin-induced sustained orofacial pain. In the early stage, the orofacial areas of the primary somatomotor cortex were separated from the other primary somatomotor cortices and integrated with subcortical and frontoparietal regions, constituting a brain-wide pain supersystem. As pain decreased over time, the subcortical and frontoparietal regions were separated from this pain supersystem and connected to multiple cerebellar regions. Machine-learning models based on these dynamic network features showed significant predictions of changes in pain experience across two independent datasets (*n* = 48 and 74). This study provides new insights into how multiple brain systems dynamically interact to construct and modulate pain experience, potentially advancing our mechanistic understanding of chronic pain.

## Introduction

The experience of pain unfolds over time and dynamically changes (Kucyi & Davis, 2015), and the sustained and spontaneously fluctuating nature is a key characteristic of clinical chronic pain (Apkarian, Krauss, Fredrickson, & Szeverenyi, 2001). Pain is known to consist of multiple component processes ranging from sensory and affective to cognitive and motivational processes (Melzack & Casey, 1968), and thus the construction and modulation of pain are subserved by distributed brain systems (Coghill, 2020; H. Mano & Seymour, 2015). Importantly, the degree to which each component process contributes to pain experience changes over time (Hashmi et al., 2013). For example, fear-avoidance (Vlaeyen & Linton, 2012), maladaptive coping (Jensen, Turner, Romano, & Karoly, 1991), and learning and memory (Apkarian, Baliki, & Geha, 2009; Phelps, Navratilova, & Porreca, 2021) become more important in explaining pain experience as pain becomes chronic. Therefore, identifying the whole-brain network features that can explain and predict the natural fluctuations and changes of sustained pain (Farmer, Baliki, & Apkarian, 2012; Kucyi & Davis, 2015) is crucial to understanding why pain naturally decreases in some cases and individuals but not in others (Ploner, Lee, Wiech, Bingel, & Tracey, 2011). In this study, we examined the dynamic reconfiguration of whole-brain functional networks underlying the natural fluctuation in sustained pain. Our key research questions include (**Figure 1A**): How does the brain community structure dynamically change (i) for the sustained pain condition compared to the resting condition and (ii) over the course of sustained pain from its initiation to remission? (iii) Can we develop predictive models of sustained pain based on the patterns of dynamic brain network changes?

**Figure 1.**
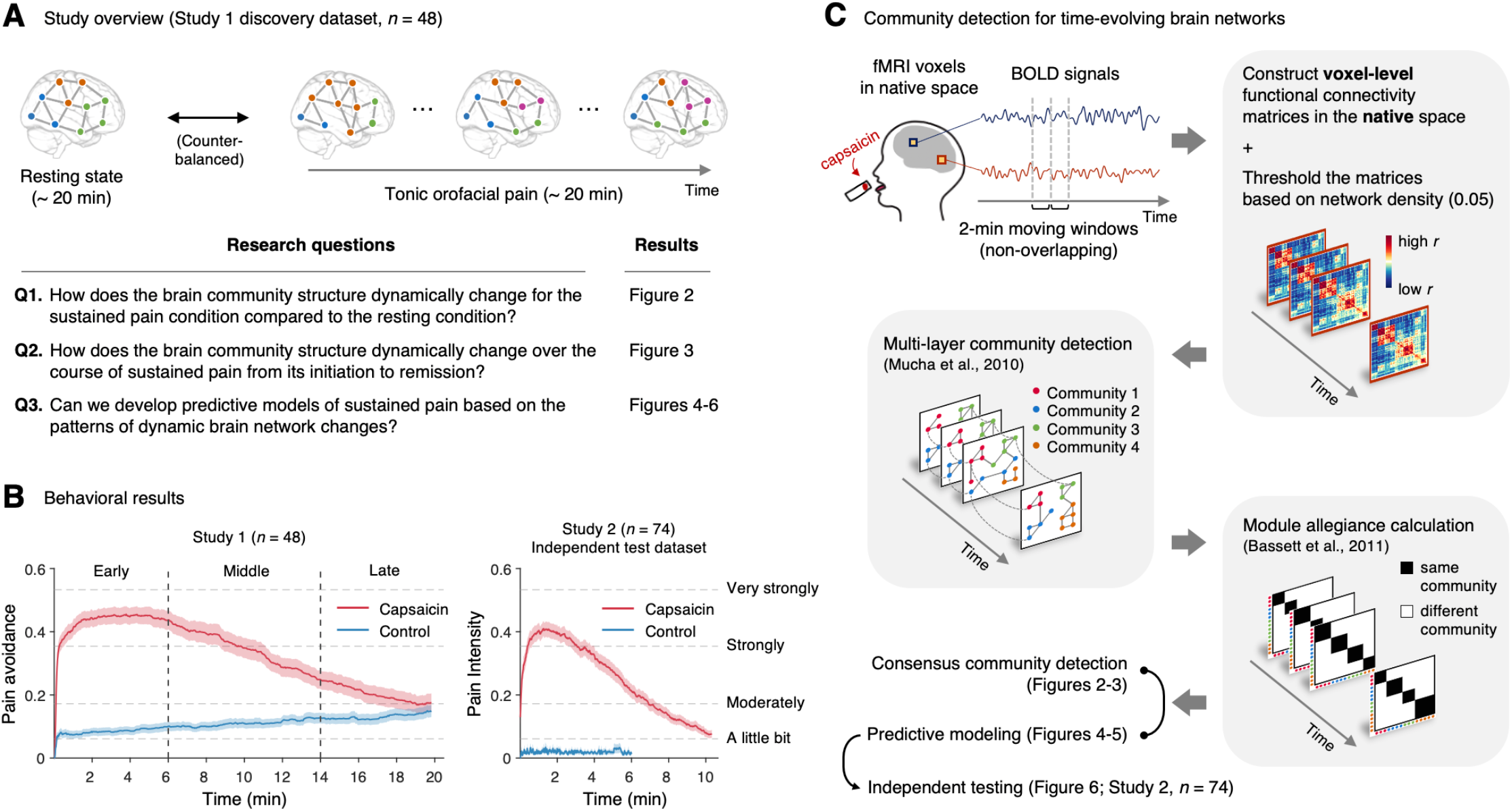
Study overview and behavioral results. **(A)** Three main research questions of the current study. We aimed to answer the research questions by examining the dynamic patterns of functional brain network reconfiguration during 20 minutes of tonic orofacial pain experience and comparing them to the pain-free resting state. **(B)** Behavioral results. We asked participants to continuously report their pain in the scanner using either a pain avoidance rating scale (“how much do you want to avoid this experience in the future?”) for Study 1 (“Discovery dataset”, *n* = 48) or a pain intensity rating scale (“how intense is your pain?”) for Study 2 (“Independent test dataset”, *n* = 74). The anchors of the scale (the horizontal dashed lines) were based on a modified version of the general Labeled Magnitude Scale (gLMS). The vertical dashed lines show how we define the early, middle, and late periods of sustained pain. The solid lines represent group mean ratings (red for capsaicin, and blue for control), and the shading represents standard errors of the mean (s.e.m.). **(C)** The overview of main analyses. Voxel-level functional connectivity was estimated in the native space using 2-min moving windows and thresholded at 0.05 network density (see Materials and Methods for details). We identified the time-evolving network community structures from these suprathreshold connectivity matrices using the multi-layer community detection method (Mucha et al., 2010), and calculated the module allegiance (Bassett et al., 2011). Using the module allegiance values as input features, we conducted predictive modeling and tested the models on Study 2 independent test dataset (*n* = 74).

To answer these questions, we conducted two functional Magnetic Resonance Imaging (fMRI) experiments with 48 and 74 healthy participants while they were experiencing 20 and 10 minutes of capsaicin-induced tonic orofacial pain, respectively. Tonic pain has long been used as an experimental model of clinical pain (Dubuisson & Dennis, 1977), and a previous study demonstrated that capsaicin-induced tonic orofacial pain showed a similar network-level neural representation as clinical pain, suggesting its potential for clinical translation (Lee et al., 2021). The application of a capsaicin-rich hot sauce on a participant’s tongue can effectively elicit pain sensation for approximately 10 minutes, followed by a slow remission of pain towards the end of the scan. This experimental paradigm is particularly suitable for identifying reliable features of the brain network for different periods of pain (i.e., initiation to remission), which has been challenging in clinical fMRI studies because of the small variance and heterogeneity of clinical pain trajectories within a few sessions of fMRI scans.

We first identified the time-evolving functional brain network structures using a community detection method for multi-layer networks (Mucha, Richardson, Macon, Porter, & Onnela, 2010) using data from Study 1 (“Discovery dataset,” *n* = 48). We conducted this network analysis at the voxel-level on an individual’s native brain space to fully utilize personalized and fine-grained pattern information of dynamic changes over time. We then compared the group-level representative brain community structures between the sustained pain condition versus the no-pain resting condition (the first research question) and between different phases of pain, including its early, middle, and late periods (the second research question). Finally, we developed machine-learning models based on the brain community patterns using the data from Study 1, either to classify pain versus no-pain conditions or to predict the dynamic changes in pain ratings, and tested the performances of these models on an unseen dataset (Study 2, “Independent test dataset,” *n* = 74) (the third research question).

The results showed that the orofacial areas (i.e., ventral part) of the primary somatomotor regions were separated from the other (i.e., dorsal) primary somatomotor regions and instead integrated with subcortical (e.g., thalamus, basal ganglia) and frontoparietal regions (e.g., dorsolateral prefrontal cortex) during sustained pain. Interestingly, this pain-induced somatomotor-dominant brain community structure, termed “pain supersystem,” (Zheng et al., 2019) changed over time. The subcortical and frontoparietal regions affiliated with the pain supersystem in the early period of pain were gradually separated from the somatomotor network and strongly connected to multiple cerebellar regions as pain decreased. Machine-learning models based on these brain network organization patterns could discriminate the sustained pain from the pain-free control condition and predict dynamic changes in pain experience including pain avoidance and pain intensity over time. The models were generalized across two independent tonic pain datasets (“Discovery dataset” and “Independent test dataset,” *n* = 48 and 74).

Overall, this study contributes to the understanding of dynamic interactions among multiple functional brain networks in response to sustained pain, offering new insights into the mechanistic understanding of chronic pain.

## Results

### Experimental design and behavioral results

In Study 1 (“Discovery dataset”), we scanned 48 participants while we delivered the capsaicin-rich hot sauce onto the participants’ tongues (“Capsaicin” condition) or had the participants rest without stimuli (“Control” condition). This experiment also included the bitter taste and aversive odor conditions, which were not analyzed here because they were not the focus of the current study. The fMRI scan duration was 20 minutes for both conditions, which was sufficient to cover the entire period of sustained pain from its initiation to the complete remission. During the scan, we asked participants to report the continuous changes in pain avoidance by asking the following question, “how much do you want to avoid this experience in the future?” With this question, we aimed to measure the continuous changes in the avoidance motivation induced by sustained pain, which is known as a core component of clinical chronic pain (Vlaeyen & Linton, 2012). We employed a modified version of the general Labeled Magnitude Scale (gLMS) to better represent pain experience at the super-high pain range (Bartoshuk et al., 2004). The order of the experimental conditions was counterbalanced across participants.

The pain avoidance ratings for the capsaicin condition were higher than those for the control condition throughout the 20 minutes of the experiment (**Figure 1B**), 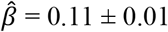 (mean ± standard error of the mean [s.e.m.]), *z* = 4.25, *P* = 2.10 × 10^−5^ (multi-level general linear model with bootstrap tests, 10,000 iterations, two-tailed, gender and the order of experimental conditions were modeled as covariates), indicating that the capsaicin stimulation effectively induced sustained pain experience and avoidance motivation. The pain avoidance ratings during the capsaicin condition exhibit an evident rise and fall. The differences in the pain avoidance between the capsaicin versus control conditions became non-significant towards the end of the scan (from 17.3 min to the end, two-tailed *P*s > 0.05, paired *t*-test, BF_01_ = 1.01-4.71), suggesting that pain subsided. A similar rise-and-fall pattern of pain ratings was observed in Study 2 with a pain intensity question, “How intense is your pain?”, except the short duration of pain because of the smaller amount of pain stimulus (i.e., hot sauce) compared to Study 1.

### Community detection of time-evolving brain networks

To examine the dynamic reconfiguration of the whole-brain functional networks, we used a multi-layer community detection approach (Mucha et al., 2010) based on the Louvain community detection algorithm (Blondel, Guillaume, Lambiotte, & Lefebvre, 2008). This approach is designed to find community structures of time-evolving networks by connecting the same nodes across different time points (**Figure 1C**) and has been successfully used in previous studies on brain network reconfigurations for learning (Bassett et al., 2011; Bassett, Yang, Wymbs, & Grafton, 2015), working memory (Braun et al., 2015; Finc et al., 2020), planning and reasoning (Pedersen, Zalesky, Omidvarnia, & Jackson, 2018), emotion (Betzel, Satterthwaite, Gold, & Bassett, 2017), pharmacologic intervention (Braun et al., 2016).

We first divided the fMRI data into 2-minute non-overlapping time windows (i.e., 10 time-bins) and estimated the functional connectivity for each time window using Pearson’s correlation for each participant and each condition. Importantly, we computed the functional connectivity at the voxel level on each participant’s native brain space to avoid an arbitrary choice of brain parcellation schemes and minimize the potential loss of information due to anatomical normalization. Also, since many of the graph analytics were developed for sparse networks (M. E. J. Newman, 2010) and voxel-level connectivity data were likely to contain many spurious correlations, we applied proportional thresholding to functional connectivity matrices. Because an arbitrary choice of the threshold can have a substantial impact on the results (van den Heuvel et al., 2017), we selected the optimal threshold level (0.05; top 5% of connections) that maximized the differences in commonly used network attributes between the capsaicin versus control conditions (**Figure S1**; for details of how we determined the optimal threshold level, see Materials and Methods). We used this threshold level for the remaining analyses.

### Differences in consensus community structure between sustained pain versus no pain (Q1)

To address the first research question, “How does the brain community structure dynamically change for the sustained pain condition compared to the no-pain resting condition?”, we compared the consensus brain community structures for the capsaicin versus control conditions. To determine the consensus community across different individuals and times, we calculated the group-level averages of module allegiance (Bassett et al., 2011). Module allegiance is a type of connectivity measure based on the community assignment pattern— defined as a matrix *T*, whose element *T*_*ij*_ is 1 when nodes *i* and *j* are assigned to the same community and 0 when assigned to different communities. To obtain the group-level averages of module allegiance, we projected the module allegiance values onto the common MNI space and then averaged them across participants and time (**Figure 2A**). Finally, we obtained the consensus community by applying a community detection algorithm to the averaged allegiance matrix (Bassett et al., 2013). For more details on the consensus community detection, see Materials and Methods.

**Figure 2.**
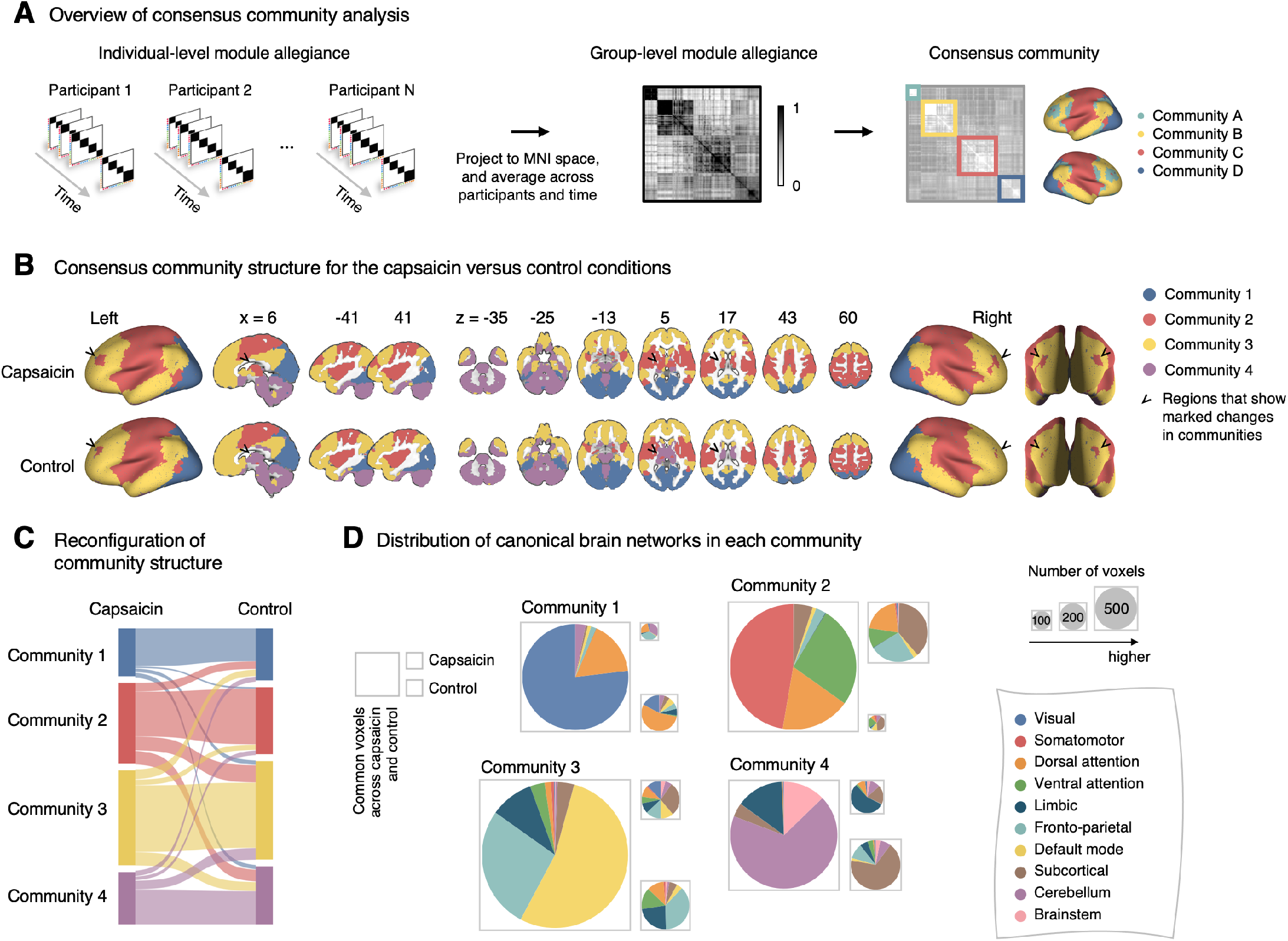
Reconfiguration of community structure for the sustained pain versus control conditions. **(A)** Analysis overview. Individual module allegiance values computed in the native spaces were projected onto MNI space and averaged across all participants and all time-bins (i.e., ten bins) to yield a group-level module allegiance matrix. Then, we identified the group-level consensus community structures by decomposing the group-level module allegiance matrix into distinct modules using the Louvain community detection algorithm (Blondel et al., 2008). **(B)** Consensus community structure for the capsaicin versus control conditions. Community colors were determined based on the canonical network membership of the largest proportion of voxels across the two conditions. V-shaped arrows indicate the regions that showed marked changes in communities. **(C)** Voxel-wise changes in community assignments between the capsaicin and control conditions. **(D)** Proportions of 10 canonical brain networks—7 resting-state large-scale networks (Yeo et al., 2011), subcortical, cerebellum, and brainstem regions—for different communities. The large square on the left shows the network composition of the voxels common across the capsaicin and control conditions, and the squares on the upper and lower right represent the voxels uniquely assigned to the community for the capsaicin or control conditions, respectively. The sizes of the squares are proportional to the number of voxels.

The consensus community structures of capsaicin and control conditions are illustrated in **Figure 2B**. We identified four main brain communities across both conditions and found the most prominent differences between the capsaicin versus control conditions in Community 2, of which the size became larger in the capsaicin condition than in the control condition (**Figure 2C**). **Figure 2D** shows the network decomposition of the communities using Yeo’s large-scale network scheme (Yeo et al., 2011)—the major player for each community was the visual network for Community 1, the somatomotor network for Community 2, the default mode network for Community 3, and the cerebellum for Community 4. The expansion of Community 2 during the capsaicin condition was mainly driven by the frontoparietal network regions from Community 3 (including the dorsolateral prefrontal and inferior parietal regions) and subcortical regions from Community 4 (including the thalamus and basal ganglia regions; **Figures 2D** and **S2**). These results suggested that sustained pain induced a global functional reconfiguration of multiple brain networks that included segregation from the default mode network and cerebellar regions and integration with the somatomotor network to form a “pain supersystem” (Zheng et al., 2019).

### Dynamic changes in consensus community structure over the course of sustained pain (Q2)

Next, to address the second research question, “How does the brain community structure dynamically change throughout sustained pain from its initiation to the remission?”, we first divided the capsaicin run into three time periods, “Early” (1-3 bins, 0-6 min), “Middle” (4-7 bins, 6-14 min), and “Late” (8-10 bins, 14-20 min). These three time periods corresponded to (1) initiation and maintenance of pain, (2) gradual pain decrease, and (3) full remission of pain, respectively (**Figure 1B**). Similar to the analysis shown in **Figure 2A**, we then averaged module allegiance matrices across participants for each period and calculated the consensus community structures from the averaged allegiance matrices.

As shown in **Figure 3**, the algorithm detected six consensus communities across time periods. The first four communities (Communities 1-4) were consistent with the communities identified in the previous analysis based on the averaged data for the capsaicin and control conditions (**Figure 2**). Patterns of dynamic changes in Communities 1-4 were similar to those from previous results. For example, Community 2 (somatomotor-dominant) was larger in the early period than in the middle and late periods (i.e., the emergence of the “pain supersystem” during the early period), and the frontoparietal network and subcortical regions were the main drivers of this change (**Figures 3A, 3C**, and **S3A**).

**Figure 3.**
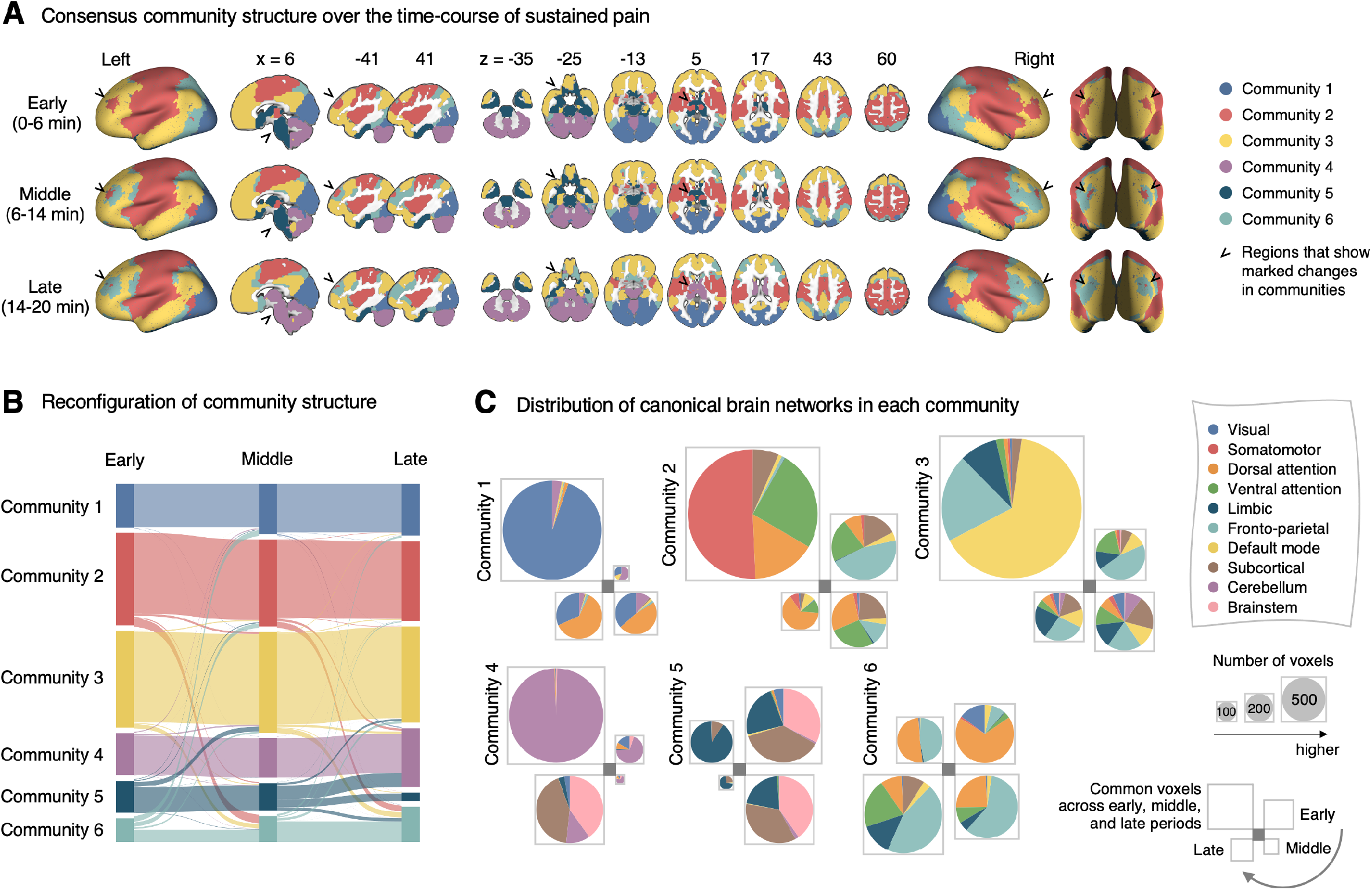
Reconfiguration of community structure over the time-course of sustained pain experience. **(A)** Consensus community structures of the early (0-6 minutes), middle (6-14 minutes), and late (14-20 minutes) periods of sustained pain. Colors indicate distinct community assignments and were determined based on the canonical network membership with the largest proportion of voxels across the periods. **(B)** Voxel-wise changes in community assignments from the early to late periods of pain. **(C)** Proportions of 10 canonical brain networks for different communities. The large square on the upper left shows the network composition of the voxels common across all periods, and the squares on the upper right, lower right, lower left represent the voxels of the early, middle, and late periods of pain after removing the common voxels, respectively. The sizes of the squares are proportional to the number of voxels.

Community 6, which was one of the newly detected communities, displayed an interesting pattern of dynamic changes over the course of sustained pain. The community mainly consisted of the frontoparietal and dorsal attention network regions and gradually increased in size over time (**Figures 3A-B**). The increase in community size in the later periods was mainly driven by the frontoparietal network regions that were among Community 2 (somatomotor-dominant) during the early period. In the later periods, these frontoparietal regions, including the dorsolateral prefrontal and inferior parietal regions, changed their affiliation to Community 6, forming a separate frontoparietal-dominant community (**Figure S3C**).

Community 5 also showed an interesting pattern. Community 5 mainly consisted of the limbic cortices and subcortical and brainstem regions during the early and middle periods. However, the subcortical and brainstem regions changed their affiliation to Community 4 (cerebellum-dominant community), leaving only a small number of voxels within the limbic regions in Community 5 in the late period (**Figures 3** and **S4A**). The composition of Community 5 during the early period suggested that the brainstem regions were closely linked with the limbic brain regions, including the medial temporal lobe structures (e.g., hippocampus, amygdala, and parahippocampal gyrus) and the temporal pole, consistent with recent findings that showed the involvement of the spino-parabrachio-amygdaloid circuit in sustained pain (Chiang et al., 2019; Huang et al., 2019; Rodriguez et al., 2017). In addition, the reconfiguration of Community 4 showed that the cerebellum-dominant community extended its connections to the brainstem and thalamus during the late period of sustained pain (**Figures 3C** and **S4B**), suggesting an important, though less investigated, pain-modulatory role of the cerebellum in pain (Claassen et al., 2020; Moulton, Schmahmann, Becerra, & Borsook, 2010).

### Module allegiance-based classifier for sustained pain versus no pain (Q3-1)

To further characterize the functional network changes induced by sustained pain, we conducted predictive modeling using the module allegiance patterns (3^rd^ research question, “Can we develop predictive models of sustained pain based on the patterns of dynamic brain network changes?”). To obtain module allegiance matrices on a common feature space and also to reduce the computational burden for model fitting, we projected a whole-brain parcellation comprising 263 regions defined on the MNI space (Schaefer atlas (Schaefer et al., 2018) with the additional brainstem, cerebellum, and subcortical regions; see Materials and Methods for details) onto an individual’s native space (**Figure 4A**). Then, we averaged module allegiance for each region, resulting in a 263 × 263 module allegiance matrix for each participant and for each time window. With the module allegiance matrices, we conducted two different types of predictive modeling: classification (developing a support vector machine [SVM] classifier to discriminate the capsaicin condition from the control condition) and regression (developing a principal component regression [PCR] model to predict the fluctuations in sustained pain ratings).

**Figure 4.**
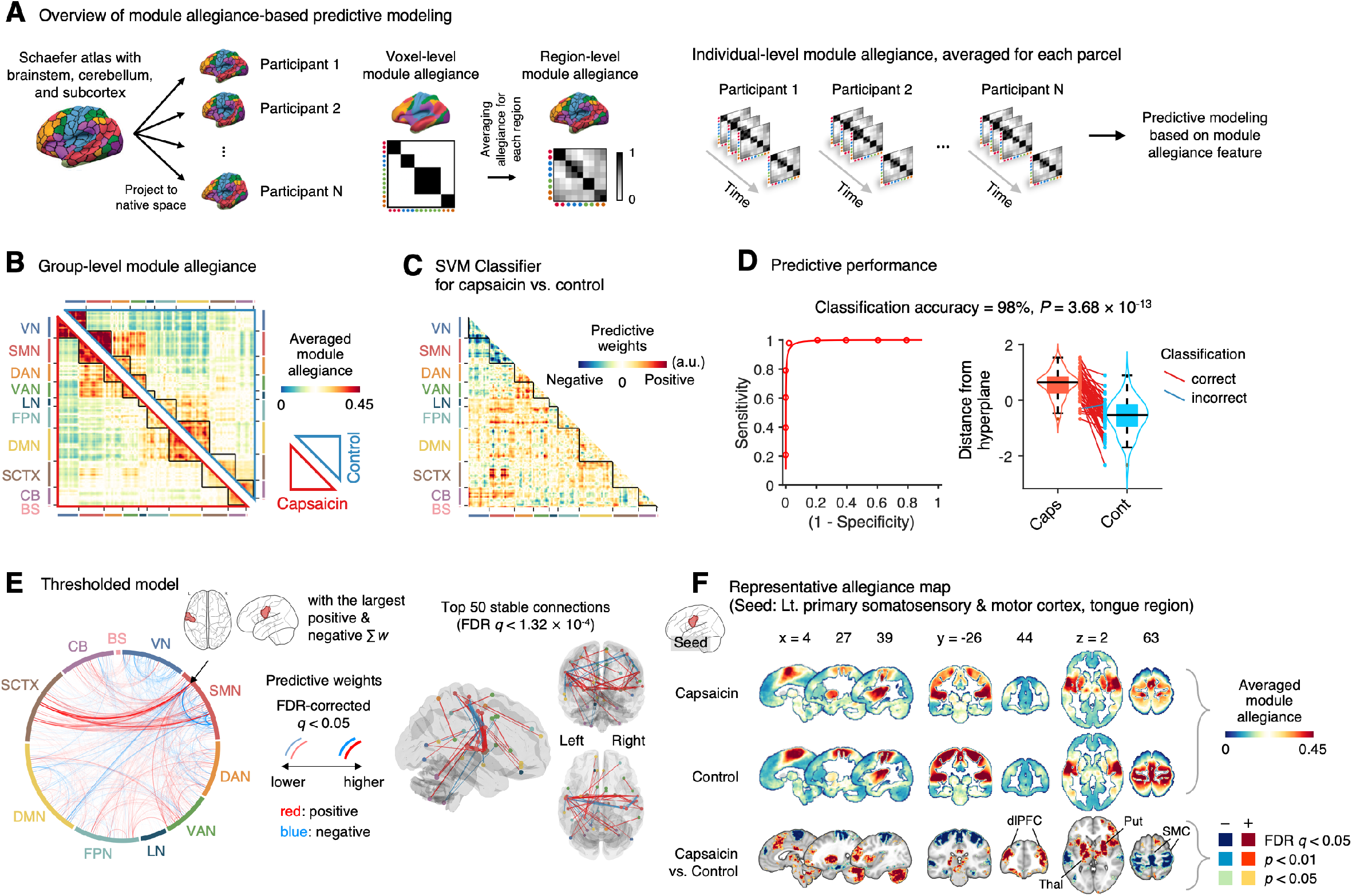
Module allegiance-based classifier for sustained pain versus resting. **(A)** The analysis overview of the module allegiance-based predictive modeling. We first projected the whole-brain atlas onto an individual’s native brain space. We then averaged the voxel-level module allegiance values for each region to create region-level module allegiance matrices. These region-level module allegiance values were used to predict whether an individual was in pain or not. **(B)** Group-level module allegiance matrix. We averaged the region-level module allegiance matrices across all participants and all time-bins, and sorted the brain regions according to their canonical functional network membership. Lower and upper triangles represent the allegiance matrices of the capsaicin and control conditions, respectively. **(C)** Raw predictive weights of the SVM classifier. **(D)** To obtain an unbiased estimate of the classifier’s classification performance, we conducted the forced-choice test with leave-one-participant-out cross-validation. Left: receiver-operating characteristics (ROC) curve. Right: cross-validated distance from hyperplane for different conditions. Each line connecting dots represents an individual participant’s paired data (red: correct classification, blue: incorrect classification). *P*-value was based on a binomial test, two-tailed. **(E)** Thresholded weights based on bootstrap tests with 10,000 iterations. Left: Predictive weights thresholded at FDR-corrected *q* < 0.05 (which corresponds to uncorrected *P* < 0.003), two-tailed. We indicated the brain region with the largest weighted degree centrality for positive and negative weights based on the thresholded model with the black arrow on the plot. Right: Top 50 stable predictive weights (FDR *q* < 1.32 × 10^−4^, uncorrected *P* < 1.91 × 10^−7^, two-tailed). **(F)** Seed-based allegiance map for the hub region that we identified in **(E)**, left primary somatosensory and motor cortex (tongue region). The bottommost row shows the contrast map for the capsaicin versus control conditions, thresholded at *t*_47_ = 2.82, FDR *q* < 0.05 (uncorrected *P* < 0.007), two-tailed, paired *t*-test. Put, putamen; dlPFC, dorsolateral prefrontal cortex; Thal, thalamus; SMC, somatomotor cortex; VN, visual network; SMN, somatomotor network; DAN, dorsal attention network; VAN, ventral attention network; LN, limbic network; FPN, frontoparietal network; DMN, default mode network; SCTX, subcortical regions; CB, cerebellum; BS, brainstem.

To develop a classifier for the capsaicin versus control conditions, we averaged the module allegiance data across time points, creating one allegiance matrix per person and experimental condition. **Figure 4B** displays the group averages of the module allegiance matrices for the capsaicin condition (lower triangle) and the control condition (upper triangle). As shown in **Figures 4C-D**, the module allegiance-based classifier showed a high classification accuracy for the capsaicin versus control conditions in a forced-choice test (accuracy = 98%, *P* = 3.68 × 10^−13^, binomial test, two-tailed; **Figure 4D**), suggesting that the individuals’ brain community structures had enough information to detect the existence of sustained pain. When we thresholded the classifier weights to examine the important features of the model based on bootstrap tests (false discovery rate [FDR]-corrected *q* < 0.05, which corresponds to uncorrected *P* < 0.003, two-tailed; the left panel of **Figure 4E**), the results suggested that the important network features of sustained pain included (1) segregation within the somatomotor network (negative weights among the brain regions within the somatomotor network in **Figure 4E**) and (2) integration between the subcortical regions and the somatomotor network regions (positive weights between the somatomotor network regions and the subcortical regions in **Figure 4E**). We could not observe the segregation within the somatomotor network in the group-level analysis presented in the previous section (**Figure 2**), suggesting that the individual-level analysis based on module allegiance provides additional insights into the dynamic network changes during sustained pain.

To further examine the important connections for the dynamic features, we visualized the top 50 stable connections (FDR *q* < 1.32 × 10^−4^, uncorrected *P* < 1.91 × 10^−7^, two-tailed) using a glass brain (the right panel of **Figure 4E** and **Table S1**). We also displayed these important connections focusing on the somatosensory and insular cortical regions. We then placed them on the sensory homunculus (**Figure S5)** to highlight that the negative weight connections were between the ventral (tongue area) and dorsal parts (other body areas) of the primary somatomotor cortex, and the positive weight connections between the tongue primary somatomotor cortex and some subcortical regions including the thalamus and basal ganglia.

We selected hub regions with the largest weighted degree centrality to provide a more detailed picture of the network reconfiguration, separately for the positive and negative weights, based on the thresholded model at FDR *q* < 0.05. The left ventral primary somatomotor region (tongue area) was selected as the sole hub region for both positive and negative weights. We then obtained a seed-based module allegiance map using the hub region as the seed. As shown in **Figure 4F**, the results reconfirmed that the hub region showed substantially decreased allegiance with the dorsal primary somatomotor regions and increased allegiance with the basal ganglia and thalamus in the capsaicin condition (thresholded at *t*_47_ = 2.82, FDR *q* < 0.05, uncorrected *P* < 0.007, paired *t*-test between capsaicin and control conditions, two-tailed). Moreover, the hub region showed increased allegiance with dorsolateral prefrontal cortex regions, consistent with previous findings that sustained pain induced integration between the frontoparietal and somatomotor networks (**Figure 2**).

### Module allegiance-based prediction model of pain ratings (Q3-2)

Next, we developed a PCR model to predict pain ratings. We used region-level module allegiance matrices across 10 time-bins of all participants as features. Because the number of features (_263_C_2_ = 34,453) was higher than the number of observations (10 ratings × 48 participants = 480), we first reduced the dimensionality of the features using principal component analysis (PCA). We then regressed the pain ratings on the principal components of module allegiance. We selected the number of principal components that yielded the best predictive performance in leave-one-subject-out cross-validation (**Figure S6**). The newly developed PCR model (**Figure 5A**) showed significantly high prediction performance (mean prediction-outcome correlation *r* = 0.29, *P* = 7.27 × 10^−6^, bootstrap test, two-tailed; mean squared error = 0.043 ± 0.006 [mean ± s.e.m.]; **Figure 5B**). Note that this model performance is biased because we conducted hyper-parameter tuning (i.e., the number of principal components) using the same dataset. Hence, we tested the model on an additional independent dataset (Study 2, *n* = 74) to provide an unbiased model performance (for the test results, see the next section).

**Figure 5.**
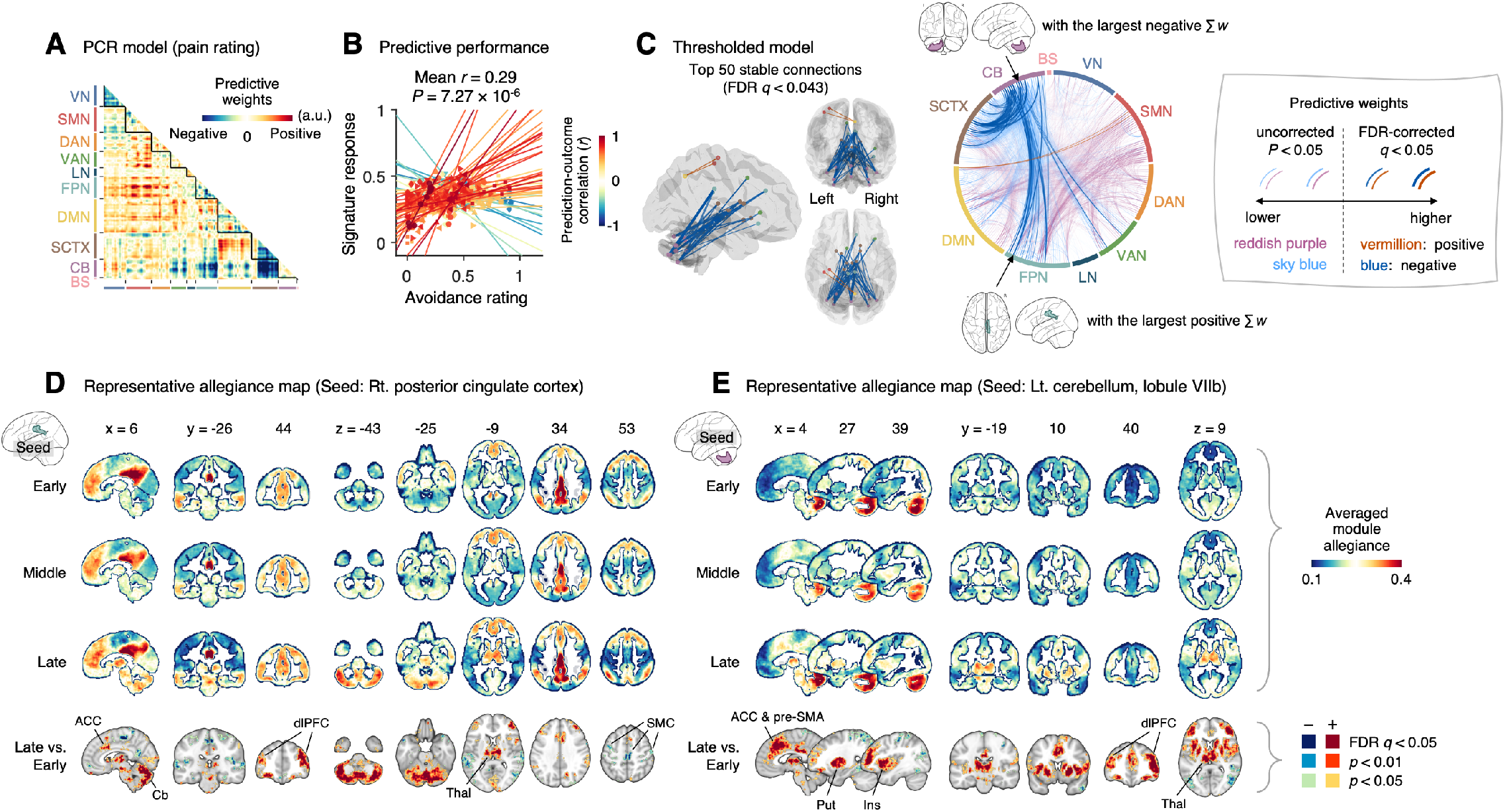
Module allegiance-based prediction model of pain rating. **(A)** The raw predictive weights of the PCR model based on region-level time-bin module allegiance matrices. **(B)** Actual versus predicted ratings. Each colored line (and symbol) represents individual participant’s ratings (10 time-bin average ratings per participant) during the capsaicin run (red: higher *r*, yellow: lower *r*, blue: *r* < 0). *P*-value is based on bootstrap tests, two-tailed. **(C)** Thresholded weights based on bootstrap tests with 10,000 iterations. Left: Top 50 stable predictive weights (FDR-corrected *q* < 0.043, which corresponds to uncorrected *P* < 6.09 × 10^−5^, two-tailed). Right: FDR *q* < 0.05 (which corresponds to uncorrected *P* < 9.24 × 10^−5^, vermillion and blue colors) or uncorrected *P* < 0.05 (reddish-purple and sky-blue), two-tailed. The brain region with the largest weighted degree centrality separately for positive and negative weights was indicated with the black arrows. **(D and E)** Seed-based allegiance maps for the hub regions across different periods of sustained pain. The right PCC and left cerebellum lobule VIIb were identified as hub regions for the positive and negative weights, respectively. The bottommost row shows the contrast map for the late versus early periods, thresholded at *t*_47_ = 3.13 **(D)** and 3.22 **(E)**, FDR *q* < 0.05 (which corresponds to uncorrected *P* < 0.003 **(D)** and 0.002 **(E)**), two-tailed, paired *t*-test. ACC, anterior cingulate cortex; Cb, cerebellum; dlPFC, dorsolateral prefrontal cortex; Thal, thalamus; SMC, somatomotor cortex; pre-SMA, pre-supplementary motor area; Put, putamen; Ins, insula; VN, visual network; SMN, somatomotor network; DAN, dorsal attention network; VAN, ventral attention network; LN, limbic network; FPN, frontoparietal network; DMN, default mode network; SCTX, subcortical regions; CB, cerebellum; BS, brainstem.

When we thresholded the predictive model based on bootstrap tests (**Figure 5C**, FDR-corrected *q* < 0.05, uncorrected *P* < 9.24 × 10^−5^, two-tailed), the results showed a dissociation between the cerebellum and the subcortical and frontoparietal regions for the high levels of sustained pain (negative weights in the circos plot in **Figure 5C**). Because few connections with positive weights were survived at the FDR correction, we examined the weight patterns at a more liberal threshold (uncorrected *P* < 0.05, two-tailed; the sky-blue and reddish-purple connections in **Figure 5D**). We observed many positive connections between the somatomotor and frontoparietal network regions, suggesting integration between the somatomotor and frontoparietal networks at high levels of sustained pain. The top 50 stable features (FDR *q* < 0.043, uncorrected *P* < 6.09 × 10^−5^, two-tailed; the left panel of **Figure 5C** and **Table S2**) were mostly negative weight connections that were connected to multiple cerebellar regions.

When we selected the hub regions with the largest weighted degree centrality based on the thresholded model at uncorrected *P* < 0.05, the right posterior cingulate cortex (PCC) within the frontoparietal network (MNI center: 6, −26, 30) and the lobule VIIb of the left cerebellum (MNI center: −26, −66, −50) were selected for the positive and negative weights, respectively. **Figures 5D** and **5E** show seed-based module allegiance maps. The right PCC region showed decreased connections with the somatomotor regions and increased connections with the dorsolateral prefrontal cortex, anterior cingulate cortex, thalamus, and cerebellar regions during the late period of pain compared to the early period (thresholded at *t*_47_ = 3.13, FDR *q* < 0.05, uncorrected *P* < 0.003, paired *t*-test, two-tailed). The left cerebellar lobule VIIb showed increased connections with the dorsolateral prefrontal cortex, anterior cingulate cortex and pre-supplementary motor area, insula, thalamus, and basal ganglia during the late period of pain compared to the early period (thresholded at *t*_47_ = 3.22, FDR *q* < 0.05, uncorrected *P* < 0.002, two-tailed). This implicates that both the right PCC and the left cerebellum may play essential roles in integrating frontoparietal and subcortical regions to construct functional communities separate from the somatomotor regions as pain decreased.

These results suggested that the frontoparietal network interacted with the somatomotor network during the early period of sustained pain. However, these connections were weakened, and the frontoparietal and subcortical regions were connected to the cerebellar regions as pain decreased.

### Testing the module allegiance-based predictive models on an independent dataset

Although our module allegiance-based predictive models demonstrated significant cross-validated prediction performances in our discovery dataset (*n* = 48), these results could be biased toward the training data. Thus, to provide unbiased estimates of model performance, we tested our models on an independent test dataset (Study 2, *n* = 74). Study 2 dataset had the same experimental design, but with a shorter scan duration—the “Capsaicin” run was 10 min, and the “Control” run was 6 min. Also, the pain rating was collected using pain intensity scale, not the pain avoidance scale, to ensure generalizability of the allegiance-based PCR model. The test results showed significant classification and prediction performances. The accuracy of the SVM model in classifying the capsaicin versus control conditions was 81%, *P* = 6.22 × 10^−8^, binomial test, two-tailed (**Figure 6A**), and the average prediction-outcome correlation of the PCR model was *r* = 0.32, *P* = 1.20 × 10^−7^, bootstrap test, two-tailed, mean squared error = 0.041 ± 0.004 (**Figure 6B**).

**Figure 6.**
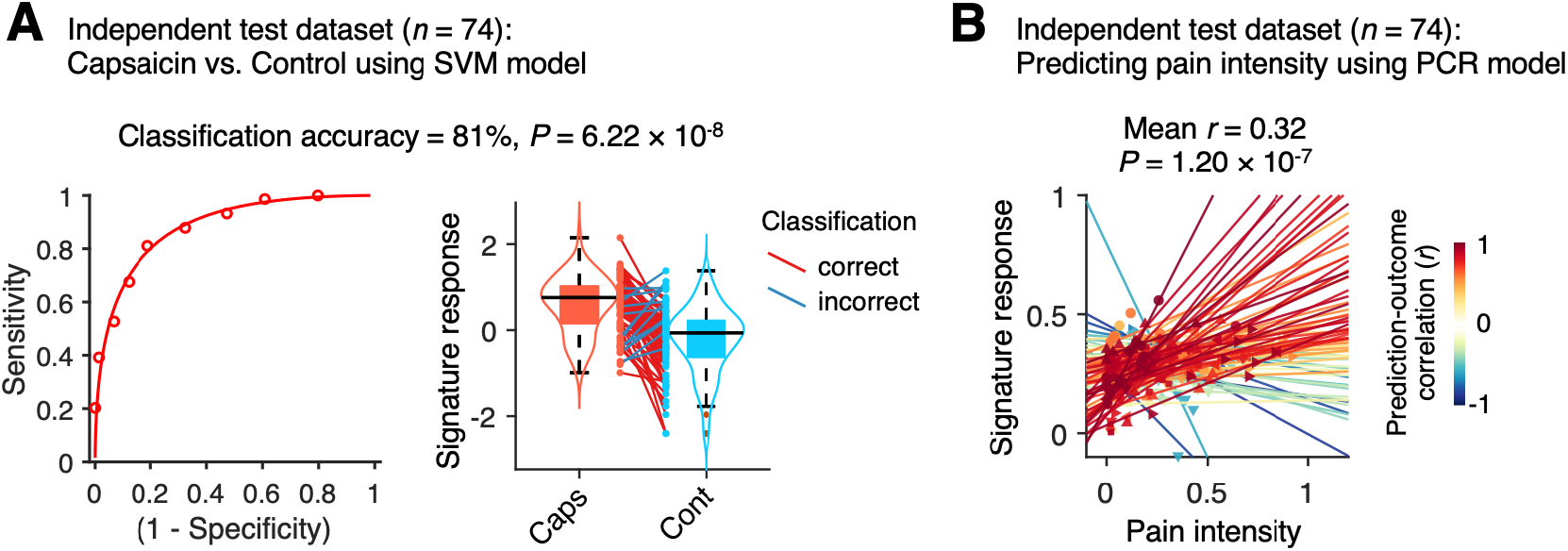
Model performance on an independent test dataset. To provide unbiased estimates of prediction performance and test the generalizability of the module allegiance-based predictive models, we tested the SVM and PCR models on an independent test dataset (Study 2, *n* = 74). **(A)** We conducted a forced-choice test to compare the signature responses for the capsaicin versus control conditions. Left: receiver-operating characteristics (ROC) curve. Right: signature response for different conditions. Each line connecting dots represents an individual participant’s paired data (red: correct classification, blue: incorrect classification). *P*-value was based on a binomial test, two-tailed. **(B)** Actual versus predicted ratings. Each colored line (and symbol) represents individual participant’s ratings during the capsaicin run (red: higher *r*, yellow: lower *r*, blue: *r* < 0). *P*-value was based on bootstrap tests, two-tailed.

## Discussion

In this study, we investigated the dynamic reconfiguration of functional brain networks during sustained pain. The main findings of this study are summarized in **Figure 7**. We compared the network structures for the capsaicin versus control conditions (**Figure 7A**) and found that (1) the somatomotor dominant community was enlarged during sustained pain by incorporating multiple subcortical and frontoparietal regions, resulting in the emergence of “pain supersystem”. (2) The ventral primary somatomotor region (tongue area) was segregated from the dorsal primary somatomotor regions and (3) integrated with the subcortical and frontoparietal regions. When we examined the dynamic network changes over time (**Figure 7B**), we observed that (4) the brainstem regions were connected to the limbic brain regions (e.g., hippocampus, amygdala, parahippocampal gyrus, and temporal pole) during the early period of pain, but soon these connections were lost as pain decreased. (5) During the late period, the frontoparietal regions were dissociated from the somatomotor-dominant community and formed their own community. Lastly, (6) a module allegiance-based predictive model of pain ratings showed that cerebellar connections with the frontoparietal and subcortical regions were important for pain decrease.

**Figure 7.**
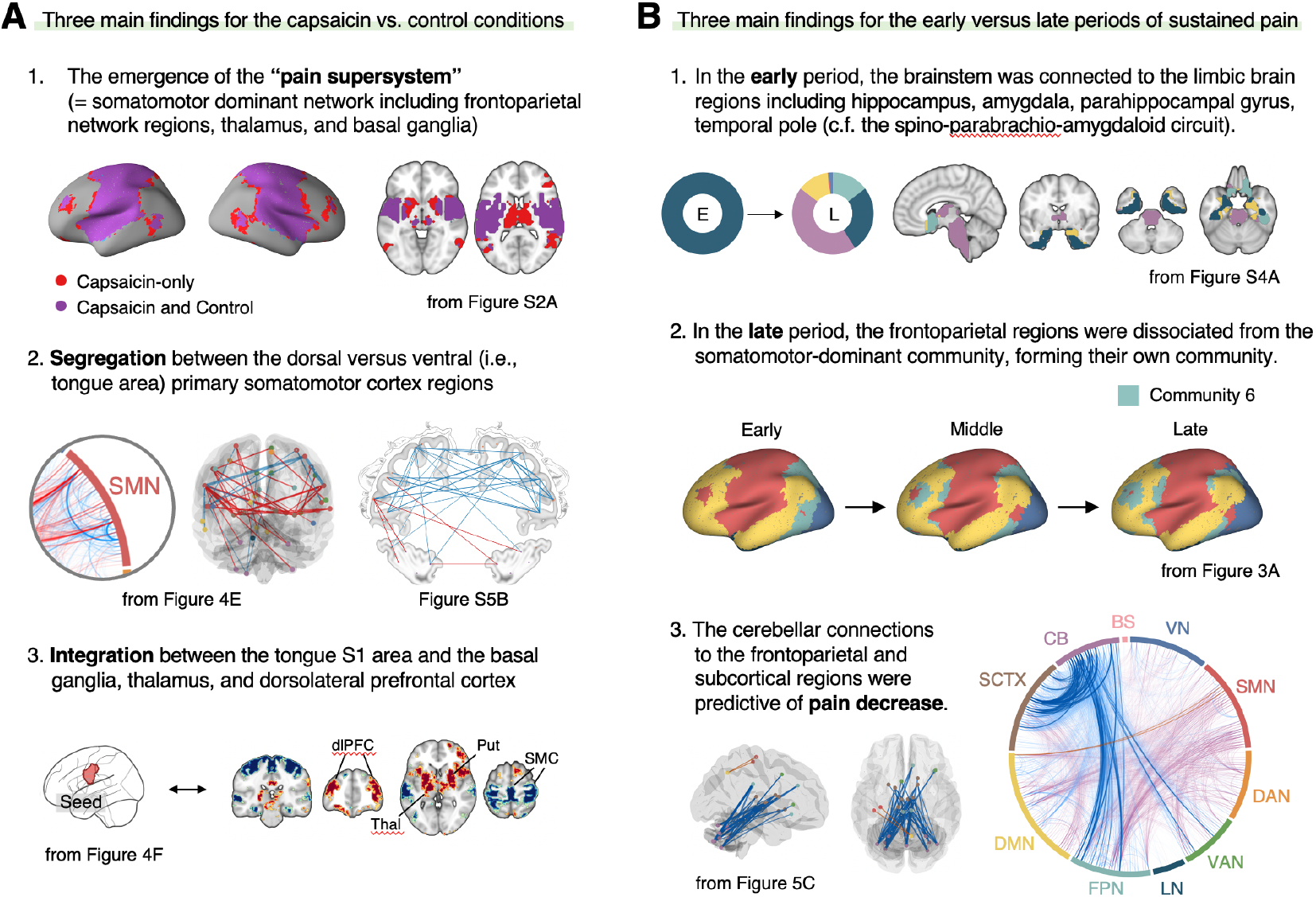
Graphical summary of the main findings. We summarized the main findings to determine the main conclusion of the current study. **(A)** Three main findings from the comparisons between the functional brain architectures of sustained pain (i.e., capsaicin) versus pain-free resting state (i.e., control) conditions (related to **Q1** and **Q3** in **Figure 1A**). **(B)** Three main findings from the comparisons between the early versus late periods of sustained pain (related to **Q2** and **Q3** in **Figure 1A**). For a detailed description of the summary, see Discussion.

Although researchers have highlighted the importance of understanding the “dynamic pain connectome” (Kucyi & Davis, 2015), previous studies were limited mainly to examining summary metrics of dynamic connectivity, such as mean and standard deviation (Bosma et al., 2018; Cheng et al., 2018; Cottam, Iwabuchi, Drabek, Reckziegel, & Auer, 2018). Therefore, a detailed characterization of how multiple brain networks dynamically interact throughout pain has yet to be conducted. A few studies examined multiple brain states over a period of pain (M. J. Lee et al., 2019; Robinson, Atlas, & Wager, 2015), but their experimental designs and analyses were not suitable to associate the changes in brain states with different phases of pain, i.e., from initiation, peak, and to remission. To better characterize the dynamic changes in functional brain community structures for different phases of pain experience, we used oral delivery of capsaicin to induce sustained and gradually decreasing pain for approximately 20 min. Although future studies need to examine whether our findings can be generalized to other stimulus modalities of sustained pain, the current study provides a reliable experimental paradigm to study time-varying network dynamics during sustained pain and spontaneous pain decrease.

One of the most critical observations in the current study is the emergence of an extended somatomotor-dominant brain community in response to sustained pain. This finding is consistent with a previous study in which subcortical and frontoparietal regions were integrated with the somatomotor network to constitute the “pain supersystem” for a brief phasic heat pain stimulus (Zheng et al., 2019). The integration of the somatomotor and frontoparietal regions can be explained using the global workspace theory (Baars, 2002), in which the conscious experience of pain requires frontoparietal involvement to enable recurrent processing of nociceptive information across multiple brain areas (Bastuji, Frot, Perchet, Magnin, & Garcia-Larrea, 2016). Also, considering that subcortical structures, such as the thalamus, are important for the early processing of nociceptive signals from the peripheral nervous system, our finding suggests that the recruitment of both bottom-up (subcortical) and top-down (frontoparietal) systems and their integration are crucial to the conscious experience of sustained pain. Previous studies that reported modular reorganization of the somatomotor network (H Mano et al., 2018) and frontoparietal network (Barroso et al., 2021; H Mano et al., 2018) in chronic pain are also in line with our results. Importantly, this systems-level integration occurred in a somatotopy-specific way—the SVM classifier (**Figure 4**) showed that the ventral primary somatomotor area, which subserves tongue sensation, was segregated from the other somatomotor regions and integrated with the subcortical regions during the early period of pain. These results are consistent with previous studies of sustained pain, which showed that the functional connectivity of the somatotopic primary somatomotor area was decreased for the other somatomotor areas, while being increased for other networks, such as the salience network (Kim et al., 2015; Kim et al., 2013; Kim et al., 2019; J. Lee et al., 2019).

In addition, we characterized how the functional brain networks were dynamically reconfigured over the period of sustained pain from the initiation to its resolution and found that the cerebellar regions played an important role in pain decrease. During the early period of pain, the subcortical and frontoparietal regions were integrated with the somatomotor network. However, as pain is resolved, these regions shifted their connections to multiple cerebellar regions. Although pain neuroimaging studies have consistently reported cerebellar activations (Moulton et al., 2010), its functional role in pain remains unclear. Interestingly, the cerebellum not only projects efferent nociceptive neurons to the thalamus (Liu, Qiao, & Dafny, 1993), but it also has afferent and efferent interconnections via the thalamus to the dorsolateral prefrontal (Middleton & Strick, 2001) and inferior parietal (Clower, West, Lynch, & Strick, 2001) cortices, which are the main components of the frontoparietal network (Allen et al., 2005; Dosenbach, Fair, Cohen, Schlaggar, & Petersen, 2008). These neuronal connections implicate the cerebellum’s functional roles in top-down pain regulation. Furthermore, recent studies suggest that cerebellum plays an important role in pain-related prediction and its error (Ernst et al., 2019), which are keys to the fear-avoidance (Vlaeyen & Linton, 2012) and motivation-decision (Fields, 2018) model of sustained pain. Therefore, our findings provide additional evidence that the cerebellar regions may be important drivers of endogenous modulation of pain avoidance and spontaneous coping responses to sustained pain. A previous lesion study in which patients with cerebellar damage exhibited increased pain sensitivity and decreased placebo analgesia (Ruscheweyh et al., 2014) supports our interpretation.

The predictive modeling based on individual-level module allegiance values (**Figures 4-5**) provided novel findings complementary to the group-level consensus community analysis (**Figures 2-3**). For example, the dissociation between the ventral versus dorsal primary somatomotor regions during sustained pain was not evident in the group-level analysis of the consensus community structure. This finding suggests that group-level averaging could obscure fine-grained information of functional anatomy (Gordon et al., 2017) with distinct regions merging together as the same network community (Braga & Buckner, 2017). In this regard, our multivariate pattern-based predictive modeling approach can provide an alternative way to study dynamic changes in functional network structures while preserving fine-grained information of individuals (Finn et al., 2015; Rosenberg, Finn, Scheinost, Constable, & Chun, 2017). Furthermore, the predictive modeling approach can provide information about the robustness and usefulness of the multi-layer community detection method by allowing us to test the prediction performance and generalizability across two independent datasets. An interesting future direction would be to examine whether the two models (i.e., SVM and PCR models) can be generalized for clinical pain fMRI datasets (Lee et al., 2021).

This study also had some limitations. First, although the prediction performances of our module allegiance-based predictive models were significant, their effect sizes were moderate (SVM model: 81% accuracy; PCR model: mean *r* = 0.32). These moderate levels of model performance may be due to information loss during a few of the multi-layer community detection analysis steps, e.g., thresholding and binarization of the connectivity matrices. For example, a previous biomarker for sustained pain based on unthresholded functional connectivity values showed better performances than the current results in a previous study (Lee et al., 2021). Considering that the understanding of dynamic pain representations in the brain is the focus of the current study, the moderate levels of model performance may be acceptable. However, future studies can still investigate how to improve the predictive performance of the predictive models based on time-varying network dynamic features. Second, the choice of two parameters for the multi-layer community detection algorithm (i.e., intra- and inter-layer coupling parameters, γ and ω) can affect the overall results (Mucha et al., 2010). Here, we followed the conventional choice (γ = 1 and ω = 1) from a previous study (Bassett et al., 2015), but the effects of these parameters on our results remain unclear. We used the predictive modeling approach and tested the model generalizability on an independent dataset to avoid excessive reverse-inference from these parameter-sensitive results alone. However, future studies will have to assess the effects of parameter choices on the overall results in a systematic way.

Overall, the current study contributes to a deeper understanding of how multiple brain systems dynamically interact in response to pain, paving the way for developing novel brain-based interventions for sustained pain. Although further studies are needed to show that our findings can be generalized to clinically relevant sustained pain conditions, we believe that this study provides new insights into the dynamic reconfiguration of functional brain networks for pain and a novel framework to investigate the neural mechanisms of sustained pain from a dynamic network perspective.

## Materials and Methods

### Participants

Study 1 (Discovery) dataset included forty-eight healthy, right-handed participants (age = 22.8 ± 2.4 [mean ± SD], 21 female) after we excluded four participants who provided avoidance rating scores higher in the control run than the capsaicin run and one participant whose brain coverage of MRI was insufficient. This dataset was included in a previous publication as an independent test dataset (as Study 3 dataset) (Lee et al., 2021). Study 2 (Independent test) dataset included seventy-four healthy, right-handed participants (age = 22.1 ± 2.4 [mean ± SD], 34 female). All participants were recruited from the Suwon area in South Korea. The institutional review board of Sungkyunkwan University approved the study. All participants provided written informed consent. The preliminary eligibility of the participants was determined through an online questionnaire. Participants with psychiatric, neurological, or systemic disorders and MRI contraindications were excluded.

### Capsaicin stimulation

In Study 1, we applied capsaicin-containing hot sauce (a food ingredient, Capsaicin Hot Sauce from Jinmifood, Inc.) to the participants’ tongues to induce tonic pain with minimal risk. To deliver the hot sauce inside the scanner, we dropped a small amount of hot sauce (0.1 ml) onto filter paper (2 cm × 6.5 cm). We spread the hot sauce in a circular format (diameter = 1 cm) on the upper 1/3 of the filter paper. While the participants were lying in the scanner, we handed the filter paper to the participants. The participants carefully placed the capsaicin side of the paper on their tongue (the participants had an opportunity to practice this procedure with a paper without capsaicin inside of the scanner). We then asked them to close their mouths. After 30 seconds, we asked them to open their mouths and put the paper on the towel on their chest. We then asked participants to keep opening their mouths and breathing only through the mouth for one min to prevent the capsaicin liquid from flowing into the throat. In this way, the liquid dried up, and the capsaicin settled down on a specific area of the tongue. After one minute, we asked the participants to close their mouths (and keep closing their mouths) while starting the fMRI scan. The participants provided their ratings using an MR-compatible trackball device when a rating scale appeared on the screen. We used this particular procedure for the following reasons: (1) It reduces the risk of coughing in the scanner, (2) can keep the pain within a tolerable range while maximizing the pain intensity, and (3) simplifies the delivery method without additional equipment.

In Study 2, we used an almost similar capsaicin delivery procedure, but there were also some differences. First, we used a smaller amount of hot sauce (0.05 ml) because the scan duration of the independent test dataset was shorter (approximately half) than the duration of the discovery dataset. Second, capsaicin was delivered in the middle of the scan. More specifically, participants held the filter paper with their hands for the first 43 seconds (13 seconds for additional removal of volumes for image stabilization) since the start of the scan, and then placed the filter paper on their tongue and closed their mouths for 20 seconds. The participants then removed the filter paper, opened their mouths for 20 seconds, and then closed their mouths. The total duration of sustained pain was approximately 10 minutes and 20 seconds.

### Experimental design

In Study 1, there were originally four experimental conditions: (i) capsaicin, (ii) bitter taste, (iii) aversive odor, and (iv) control, but in the current study, only the capsaicin and control runs were used. For the experimental procedure of other conditions, please see the previous publication that used this dataset (Lee et al., 2021). After a structural scan, we administered the four condition runs, and the order of the conditions was counterbalanced across participants. Each run lasted for 20 minutes, and participants provided the avoidance ratings continuously throughout the run. We designed the experiment to have long scans to capture the full rise and fall of each sensation. For the rating scale, we used a modified version of the general Labeled Magnitude Scale (gLMS) (Bartoshuk et al., 2004): The anchors began with “Not at all” [0] to the far left of the scale and continued to the right in a graded fashion with anchors of “A little bit” [0.061], “Moderately” [0.172], “Strongly” [0.354], and “Very strongly” [0.533], until “Most (I never want to experience this again in my life)” [1] on the far right. To prevent participants from falling asleep and to help maintain a certain level of alertness during the scan, we used an intermittent simple response task, in which the color of the rating bar on the screen was changed from orange to red for 1 second every minute with a jitter, and the participants had to respond to the color change by clicking the button on the trackball device. During the preprocessing of the data, we included additional regressors of the color changes and button clicks to remove confounding effects related to the task. After the scan, we asked the participants multiple post-scan questions regarding their thought contents during the scan, which were not included in the current study.

Study 2 had three experimental conditions: (i) capsaicin, (ii) control, and (iii) phasic heat stimulation. We used only the capsaicin and control runs. The control and capsaicin runs were conducted at the start and the end of the session, respectively. The control run lasted for 6 minutes and 13 seconds, and the capsaicin run lasted for 11 minutes and 43 seconds. After the first 13 seconds (13-second data were discarded before further analyses), participants provided intensity ratings continuously throughout the run using a trackball. After the scan, we asked the participants multiple post-scan questions regarding their thought contents during the scan.

### fMRI data acquisition

Whole-brain fMRI data were acquired using a 3T Siemens Prisma scanner at Sungkyunkwan University. High-resolution T1-weighted structural images were also acquired. Functional EPI images were acquired with TR = 460 ms, TE = 29.0 ms, multiband acceleration factor = 8, field of view = 248 mm, 82×82 matrix, 3×3×3 mm^3^ voxels, 56 interleaved slices. Stimulus presentation and behavioral data acquisition were controlled using Matlab (Mathworks) and Psychtoolbox (http://psychtoolbox.org/).

### fMRI data preprocessing

Structural and functional MRI data were preprocessed using our in-house preprocessing pipeline (https://github.com/cocoanlab/surface_preprocessing) based on AFNI, FSL, and Freesurfer. This is similar to the Human Connectome Project (HCP) preprocessing pipeline (Glasser et al., 2013). For structural T1-weighted images, magnetic field bias was corrected, and non-brain tissues were removed using Freesurfer. Non-linear transformation parameters projecting native T1 space to MNI 2×2×2 mm^3^ template space were also calculated using FSL. For functional EPI images, the initial volumes (22 images [10 seconds] for Study 1, 29 images [13 seconds] for Study 2) of fMRI images were removed to allow for image intensity stabilization. Then, the images were motion-corrected using AFNI and distortion-corrected using FSL. These EPI images were co-registered to T1-weighted images using the boundary-based registration (BBR) technique (Greve & Fischl, 2009) that used FSL for the first registration and Freesurfer for refinement, similar to the HCP pipeline. We then removed motion-related signals from co-registered EPI images using ICA-AROMA (Pruim et al., 2015). Additional preprocessing modules, including (i) removal of nuisance covariates, (ii) linear de-trending, and (iii) low-pass filtering at 0.1 Hz, were combined and conducted in one step using the 3dTproject function in AFNI to avoid introducing unwanted artifacts (Lindquist, Geuter, Wager, & Caffo, 2019). We included mean BOLD signals from white matter (WM) and cerebrospinal fluid (CSF) (Pruim et al., 2015), and time-points where intermittent arousal maintenance tasks appeared (total of 20 times) as nuisance covariates. For computational efficiency in further analyses, including community detection, these de-noised EPI images were resampled to 4×4×4 mm^3^ spatial resolution and masked with an individually defined gray matter boundary image. This gray matter mask obtained using Freesurfer was dilated and eroded five times to create smooth edges, and then resampled to 4×4×4 mm^3^ spatial resolution to match the spatial dimension of EPI images.

### Functional connectivity calculation and proportional thresholding

Whole-brain voxel-wise functional connectivity was computed using Pearson’s correlation. To determine the optimal threshold level, we tested multiple network density thresholding options (0.01, 0.05, 0.10, 0.20, 0.30, and 0.40) and compared the global-level network characteristics between the capsaicin versus control conditions. Five network attributes, including assortativity, transitivity, characteristic path, global efficiency, and modularity, were used for comparison. We calculated the network attributes using the Brain Connectivity Toolbox (https://sites.google.com/site/bctnet/) (Rubinov & Sporns, 2010).

A brief description of these measures is as follows: *(1) Assortativity:* This attribute is related to how often each node (here, a voxel) is connected to the other nodes that have a similar number of links (here, functional connectivity) (M. E. Newman, 2002). It can be measured using Pearson’s correlation between the degrees of every pair of connected nodes. High assortativity means that there are mutual connections between high-degree hub nodes, reflecting the overall resilience of a network. The ‘assortativity_bin’ function of the toolbox was used. *(2) Transitivity:* This attribute is related to how often the two nodes that are connected to the same node are also connected to each other (Mark EJ Newman, 2003) and measured as the ratio of the number of inter-connected triplets of nodes to the number of all triplets of nodes. High transitivity indicates that nodes of a network are more likely to be clustered together. The ‘transitivity_bu’ function is used. *(3) Characteristic path:* This attribute is the average of the shortest path between all pairs of connected nodes (estimated using ‘distance_bin’ function), reflecting the functional dissociation of a network. A high characteristic path length suggests that the network is in a disintegrated state. *(4) Global efficiency:* This attribute is the average of the inverse of the shortest path between all pairs of connected nodes (estimated from ‘distance_bin’ function), reflecting the functional integration of a network. A high global efficiency suggests that a network is in an integrated state. *(5) Modularity:* This attribute is related to how much a network can be clearly divided into a set of modular structures. A network with fewer between-module connections and more within-module connections has a higher level of modularity. The Louvain community detection algorithm (Blondel et al., 2008) was used to determine the inherent community structure of a network (‘community_louvain’ function).

Using these global-level network attributes, we calculated the group-level *z*-statistics of the differences between conditions for different network density levels. Then, we selected the density level that maximized the sum of absolute *z*-scores as the optimal threshold level. As shown in **Figure S1**, we selected a network density of 0.05 (i.e., only the top 5% connections were survived) as the optimal threshold level.

### Dynamic functional connectivity (moving time window)

Whole-brain voxel-wise connectivity was computed for every 2-min non-overlapping time window within individuals using Pearson’s correlation. Proportional thresholding at a pre-determined network density of was applied to each of these dynamic connectivity matrices (Study 1 dataset: 10 matrices per run and participant; Study 2 dataset: 5 matrices per participant for the capsaicin run, and 3 matrices per participant for the control run), without binarization of connectivity values.

### Multi-layer community detectio

To identify the time-evolving network community structure, we used the multi-layer community detection approach (Mucha et al., 2010), which is a generalized version of the Louvain algorithm (Blondel et al., 2008). This method finds the optimal community structure that maximizes the ‘multi-layer modularity’ function %, which is defined as:

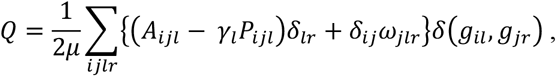

where *A*_*ijl*_ is the adjacency matrix of layer *l* (i.e., dynamic connectivity matrices), *P*_*ijl*_ is the optimization null model of layer *l* (Newman-Girvan null model 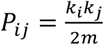, where *k*_*i =*_ *Σ*_*j*_ *A*_*ijl*_ and 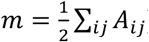 is the intra-layer resolution parameter of layer *l, ω*_*jlr*_ is the inter-layer coupling parameter between node *j* in layer *l* and node *j* in layer *r*, µ is the total sum of edge weights 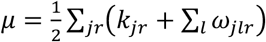 the Kronecker delta *δ*(*g*_*il*_, *g*_*jr*_)is equal to 1 when the community assignment of node *j* in layer *l* (*g*_*il*_) is the same as the community assignment of node *j* in layer *r* (*g*_*jr*_), and equal to 0 otherwise. Here, we set all the γ _*l*_ and ω _*jlr*_ parameters to 1 as in previous studies (Bassett et al., 2015).

This modularity-maximizing community detection method is inherently nondeterministic (NP-hard problem). Therefore, we iterated this procedure 100 times to determine the consensus community structure, as described in previous studies (Bassett et al., 2013; Bassett et al., 2015). Details of the within-individual consensus community detection procedure are as follows:

i. Module allegiance, which is defined as a matrix *T* whose element *T*_*ij*_ is equal to 1 if nodes *i* and node *j* are assigned to the same community and 0 otherwise, was calculated for each iteration of multi-layer community detection.
ii. The module allegiance matrices from (i) were averaged across all iterations.
iii. Null models of module allegiance matrices were obtained by repeating (i) and (ii) with a random permutation of the original community assignments.
iv. The averaged allegiance matrix from (ii) was thresholded at the maximum value of the null-model allegiance matrix from (iii).
v. The Louvain community detection algorithm was applied to the thresholded allegiance matrix with 100 iterations. If all the 100 community assignment vectors were identical, the community assignment was determined to be the consensus community. Otherwise, steps (i) to (v) were repeated, using the 100 community assignment vectors as the input of (i).

Using this within-individual consensus community detection procedure, we obtained a consensus community for each layer (i.e., 2-minute time window), run, and participant.

### Projecting individuals’ community assignments into MNI space

For analyses requiring group-level inferences, we computed one-to-one mapping between the voxels in an individual’s native space and the voxels in the MNI space as follows: First, every voxel within the resampled gray matter mask (4×4×4 mm^3^ native space; obtained from the preprocessing steps) was labeled with unique indices. Next, the labeled gray matter mask was resampled to 2×2×2 mm^3^ native space, projected to the MNI space using the pre-computed parameters for nonlinear transformation, and then resampled to 4×4×4 mm^3^ space, with the nearest neighbor interpolation method. The output contains voxel labels indicating the original location in the native space, which then can be used as a mapping rule for projecting community assignments defined in the native space onto the MNI space.

### Group-level consensus community detection (Figures 2 and 3)

To examine the group-level consensus community structure, we performed the following consensus community detection procedure:

i. The individual-level consensus community assignments in the native space were projected onto the MNI space using the pre-computed one-to-one mapping rule.
ii. The projected module community assignments were converted into module allegiance matrices.
iii. The module allegiance matrices were averaged across all individuals and across time depending on the type of consensus community. For the analyses on the capsaicin versus control conditions (as in **Figure 2**), allegiance values were averaged across all time-bins for each run. For the analyses on the early/middle/late periods (as in **Figure 3**), the allegiance values of the capsaicin run were averaged within three separate time-bins; i.e., averaging 1-3, 4-7, and 8-10 layers into early, middle, and late-period module allegiance matrices, respectively.
iv. Null models of module allegiance were obtained by repeating (i) to (iii) with a random permutation of the original community assignments.
v. The averaged allegiance matrices from (iii) were thresholded at the maximum value of the null-model allegiance matrix from (iv).
vi. The Louvain community detection algorithm was applied to the thresholded allegiance matrix with 100 iterations. If all the 100 community assignment vectors were identical, the community assignment was determined to be the consensus community. Otherwise, steps (i) to (vi) were repeated using the 100 community assignment vectors as the input of (i).

We excluded voxels that were disconnected after the iterative consensus community detection procedure or that were assigned to a community with a small number of voxels, i.e., < 20 voxels.

### Module allegiance-based predictive modeling (Figures 4 and 5)

To quantify the relative contribution of network communities to sustained pain experience, we conducted predictive modeling using module allegiance values as input features. We first projected the whole-brain atlas onto each individual’s native brain space. The whole-brain atlas originally consisted of 265 regions, but we had to exclude two cerebellar regions (vermis VIIb and X) because these regions had a small number of voxels (23 and 42 in 2×2×2 mm^3^ space, respectively) and thus were not successfully transformed to native space in a few participants, resulting in a total of 263 regions. This atlas included 200 cortical regions from the Schaefer atlas (Schaefer et al., 2018), 61 subcortical and cerebellar regions from the Brainnetome atlas (Fan et al., 2016), and the periaqueductal gray and brainstem regions used in previous studies (Beissner, Schumann, Brunn, Eisentrager, & Bar, 2014; Roy et al., 2014). Then, voxel-wise module allegiance matrices were obtained from the individual’s consensus community structure and grouped and averaged into 263×263 region-wise allegiance matrices.

For the classification problem, as shown in **Figure 4**, we averaged ten time-varying region-wise module allegiance matrices for each run (capsaicin and control runs). Then we trained a SVM classifier to determine whether a participant was in pain or not (i.e., capsaicin versus control conditions) using region-level allegiance matrices across participants (i.e., 34,453 allegiances × 2 runs × 48 participants). To obtain unbiased estimates of classification performance, we used leave-one-subject-out cross-validation on the training dataset (Study 1) and tested the model on an independent test dataset (Study 2). To quantify the model performance, we conducted a forced two-choice classification test, which directly compared the predicted values (here, distances from the hyperplane) of two conditions for each individual. This test did not require the assumption that all participants’ brain responses to stimuli are on the same scale (Wager et al., 2013).

For regression-based modeling, as shown in **Figure 5**, we trained a PCR model (Hastie, Tibshirani, & Friedman, 2009) to predict pain avoidance ratings (10 time-bins × 48 participants) based on concatenated module allegiance matrices of capsaicin run across ten time-bins and participants (i.e., 34,453 allegiances × 10 time-bins × 48 participants). We selected 14 principal components (PCs) for the regression modeling because the PC number yielded the best predictive performance (**Figure S6**). Similar to the classification model, we tested the prediction model on an independent test dataset (Study 2).

To identify important features for the classification and prediction models, we used bootstrap tests. We randomly sampled 48 participants 10,000 times with replacement and conducted model training with the resampled data. We calculated the statistical significance of predictive weights using *z*-statistics based on 10,000 samples of 34,453 predictive weights.

### Seed-based allegiance analysis (Figures 4F, 5D, and 5E)

Using the voxel-wise module allegiance in the native space, we calculated the individual’s seed-based module allegiance map, which consisted of averaged module allegiance values between voxels within a seed region and the rest of the brain. This seed-based allegiance map was then transformed into MNI space using the pre-computed one-to-one mapping between the native and MNI spaces and averaged across participants. We then conducted a paired t-test between the capsaicin versus control conditions for the classification analysis (**Figure 4F**) and between the late versus early period of pain for the regression analysis (**Figures 5D-E**).

### Signature response calculation (Figure 6)

To test the allegiance-based classification and prediction models, we calculated a signature response score (the intensity of pattern expression) using a dot product of vectorized functional connectivity with model weights.

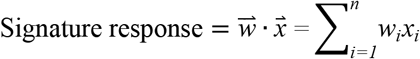

where *n* is the number of connections within the connectivity-based predictive models, *w* is the connection-level predictive weights, and *x* is the test data.

### Statistical analysis

In **Figures 4D** and **6A**, we used binomial tests for significance testing of whether forced two-choice classification accuracies were significantly higher than the probability of chance (here, 50%). The sample sizes for **Figures 4D** and **6A** were *n* = 48 and 74, respectively. In **Figures 5B** and **6B**, we conducted bootstrap tests with 10,000 iterations to test whether the sampling distribution of the within-individual prediction-outcome correlations was significantly higher than zero. Note that we performed the *r*-to-*z* transformation before the bootstrap tests. The sample sizes for **Figures 5B** and **6B** were *n* = 48 and 74, respectively. In **Figures 4E** and **5C**, we used bootstrap tests (with 10,000 iterations) to threshold each model’s predictive weights. In **Figures 4F** and **5D-E**, we used paired *t*-tests either between the seed-based allegiance maps of the capsaicin versus control conditions (**Figure 4F**) or between the seed-based allegiance maps of the late versus the early period of pain (**Figures 5D-E**). Further details of the statistical analyses are provided in each relevant description in the main manuscript.

## Supplementary Information

Figures S1-6

Tables S1-2

## Acknowledgments

We thank Hongji Kim and Soo Ahn Lee for help with conducting experiments. This work was supported by IBS-R015-D1 (Institute for Basic Science; to C.-W.W.), 2019R1C1C1004512 (National Research Foundation of Korea; to C.-W.W.), 2E30410-20-085 (KIST Institutional Program; to C.-W.W.), and by 2018H1A2A1059844 (National Research Foundation of Korea; to J.-J.L.).

## Author Contributions

J.-J.L. and C.-W.W. conceived and designed the experiment, analyzed the data, interpreted the results, and wrote the manuscript. C.-W.W. edited the manuscript and provided the supervision. S.L., D.H.L., and C.-W.W. contributed to the independent test dataset.

## Data Availability

The brain community affiliations and predictive models will be shared upon publication through a GitHub repository. The data not used in the main figures will be shared upon request.

## Code Availability

The code for generating the main figures will be shared upon publication through a GitHub repository. In-house Matlab codes for fMRI data analyses are available at https://github.com/canlab/CanlabCore and https://github.com/cocoanlab/cocoanCORE.

