## Supplementary Information for "Dynamic Functional Brain Reconfiguration During Sustained Pain"

**This PDF file includes:**

1. Figures S1-6

2. Tables S1-2

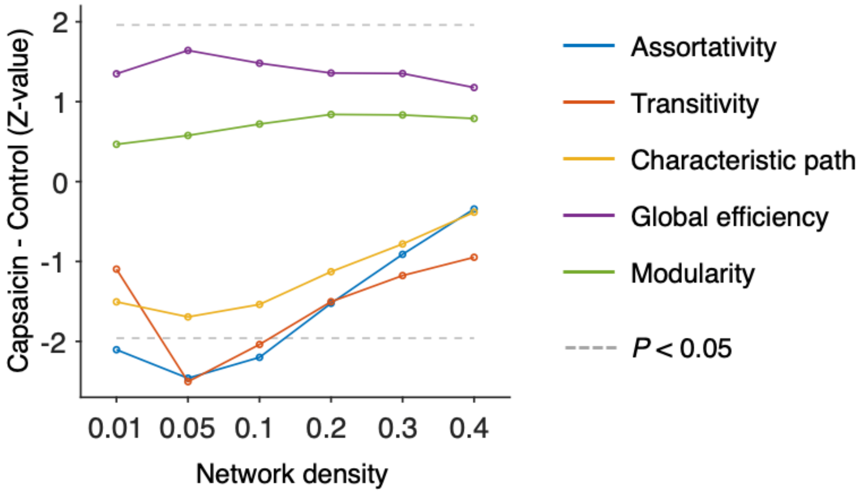


**Figure S1. Global-level network attributes.**

We compared the five global-level network attributes (i.e., assortativity, transitivity, characteristics path, global efficiency, and modularity) of the capsaicin and control conditions (*z*-values from paired *z*-test) across different levels of network density. Note that the overall differences of network attributes between the capsaicin versus control conditions were maximal at the network density of 0.05. All network attributes were measured from the binarized static connectivity matrices (for details, please see Materials and Methods).

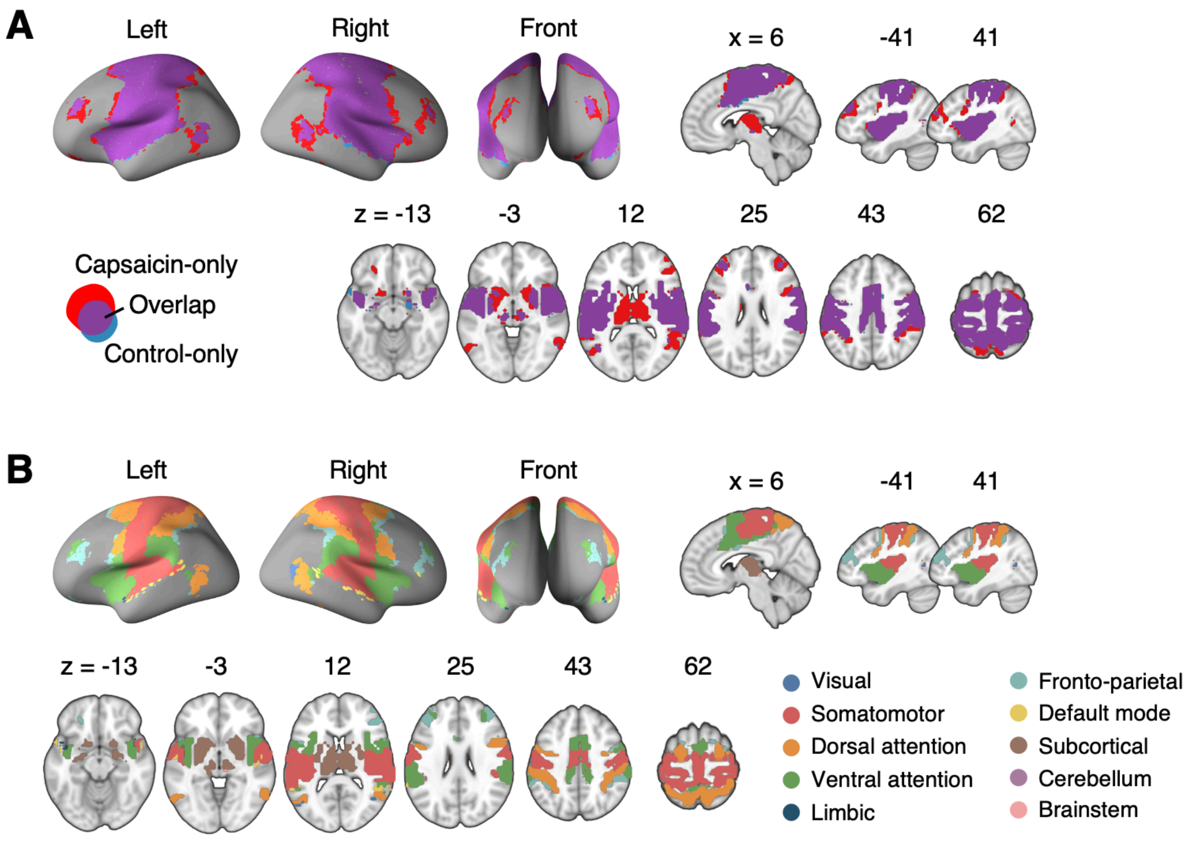


**Figure S2. Reconfiguration of the consensus Community 2.**

**(A)** We compared the spatial distributions of the consensus Community 2 (i.e., the somatomotor dominant community) of the capsaicin and control conditions. The purple regions show the overlap between the two conditions, and the red and blue regions show the unique regions for the capsaicin and control conditions, respectively. Blue region was comparatively small, indicating that the somatomotor community mainly expanded primarily during the capsaicin condition compared to the control condition.

**(B)** The spatial distribution of the ten canonical brain networks within the consensus community 2. The expansion of the Community 2 during the capsaicin condition (red regions in [**A**]) was mainly driven by the brain voxels within the frontoparietal network (e.g., dorsolateral prefrontal cortex and inferior parietal cortex) and the subcortical regions (e.g., thalamus and basal ganglia).

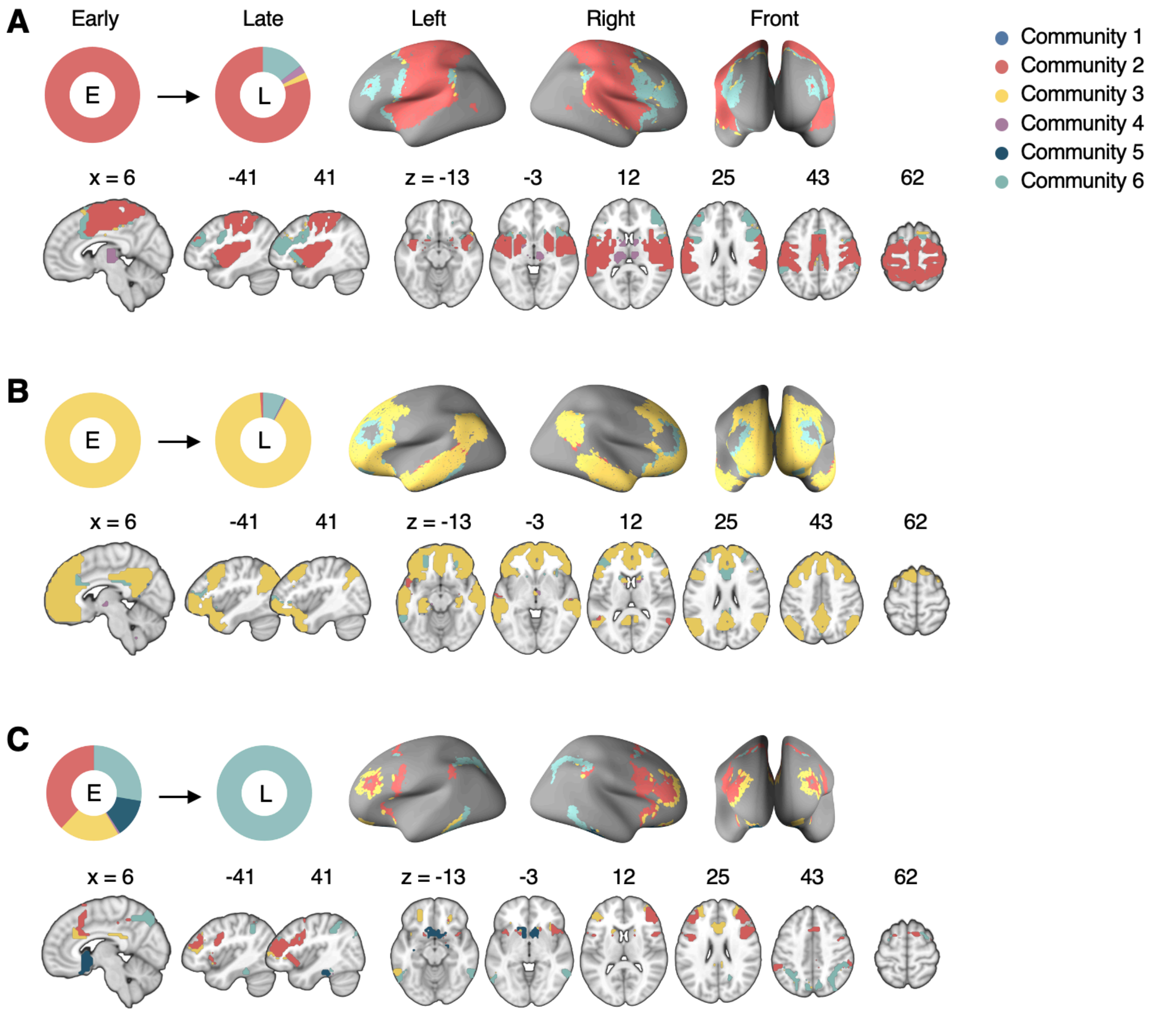


**Figure S3. Reconfiguration of the consensus Communities 2, 3, and 6.**

The reconfiguration pattern of the community assignments of the brain voxels that were assigned to **(A)** the consensus Community 2 (somatomotor network dominant community) in the early period of sustained pain, **(B)** the consensus Community 3 (default-mode network dominant community) in the early period of sustained pain, and **(C)** the consensus Community 6 (frontoparietal network dominant community) in the late period of sustained pain.

E: Early, L: Late.

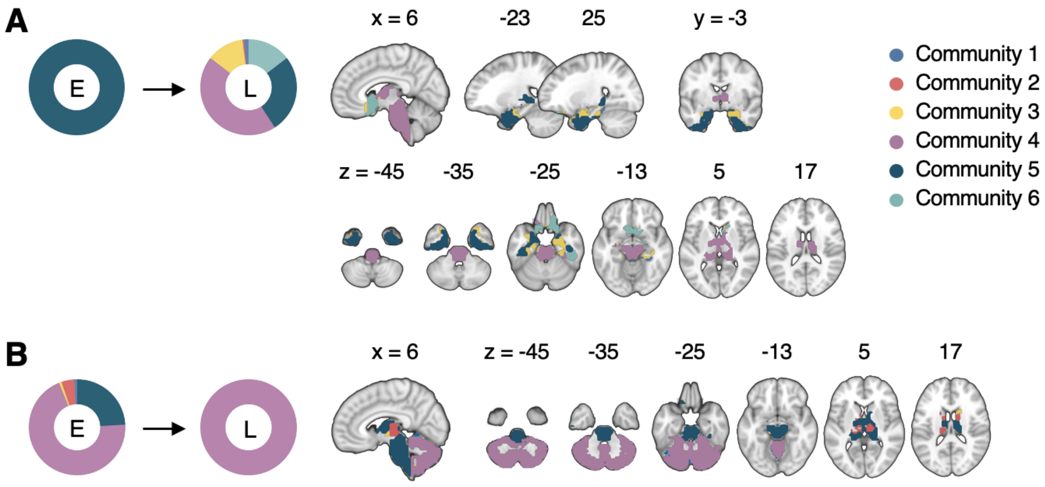


**Figure S4. Reconfiguration of the consensus Communities 4 and 5.**

The reconfiguration pattern of the community assignments of the brain voxels that were assigned to **(A)** the consensus Community 5 (limbic network dominant community) in the early period of sustained pain, and **(B)** the consensus Community 4 (cerebellum-dominant community) in the late period of sustained pain.

E: Early, L: Late.

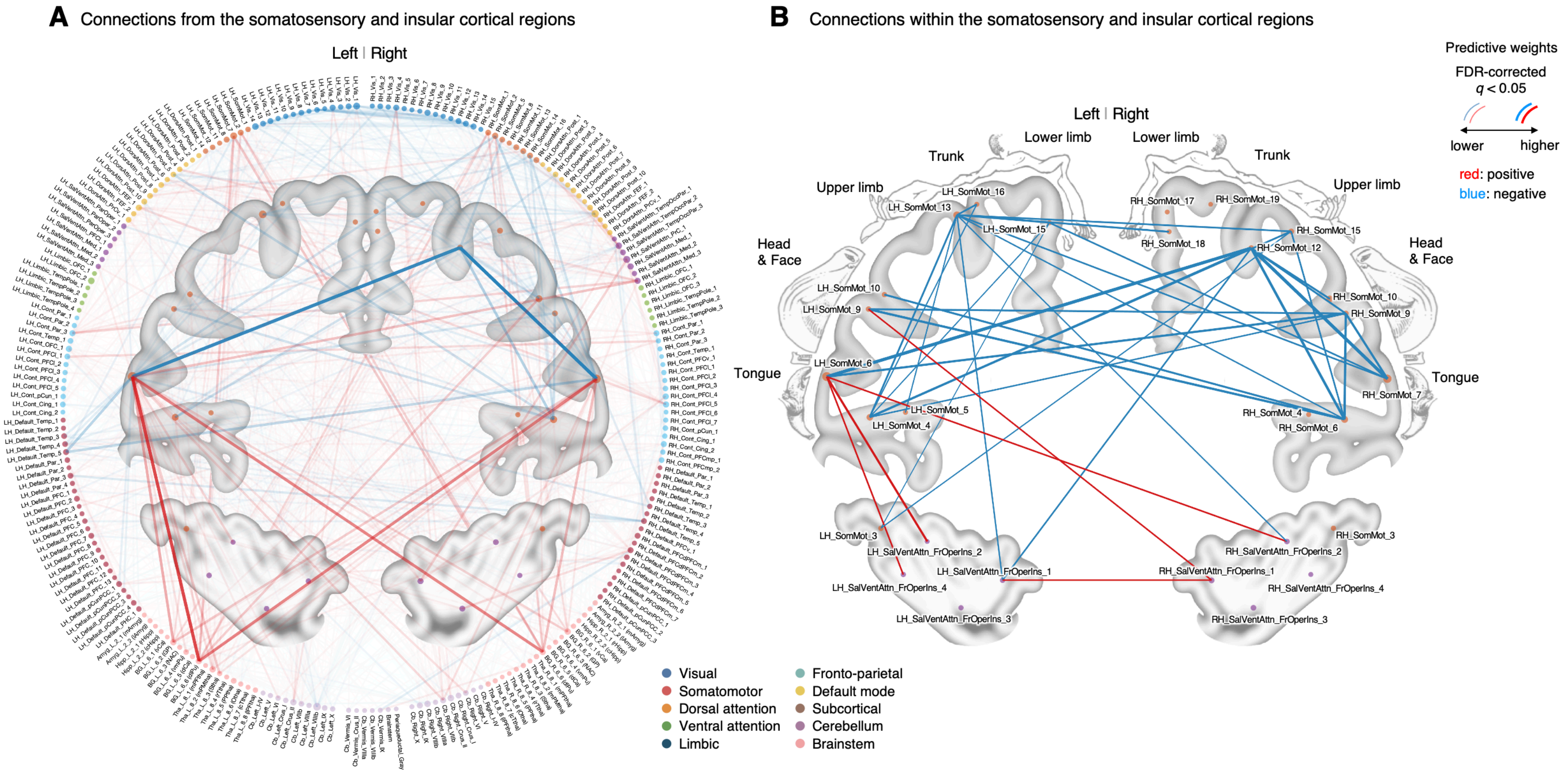


**Figure S5. Classifier weights of the somatosensory and insular cortical regions.**

**(A)** Thresholded connections (FDR $q < 0.05$, which corresponds to uncorrected $P < 0.003$, two-tailed, bootstrap test with 10,000 iterations) showing the predictive weights between the somatosensory and insular cortical regions and the other remaining whole brain regions. Line thickness and transparency indicate the absolute magnitude of predictive weights. There were strong negative weights within the somatosensory cortical regions and strong positive weights between the tongue primary somatosensory regions and subcortical regions (e.g., basal ganglia).

**(B)** Thresholded connections (FDR $q < 0.05$, which corresponds to uncorrected $P < 0.003$, two-tailed, bootstrap test with 10,000 iterations) showing the predictive weights between the primary and secondary somatosensory and insular cortical regions. Line thickness indicates the absolute magnitude of the predictive weights. Note that the location of the nodes on the brain map may not reflect the exact center coordinates of the regions-of-interests, though we marked them on the closest locations on the map. The somatosensory homunculus (modified from (Penfield & Rasmussen, 1950)) represents the overall somatotopic gradients.

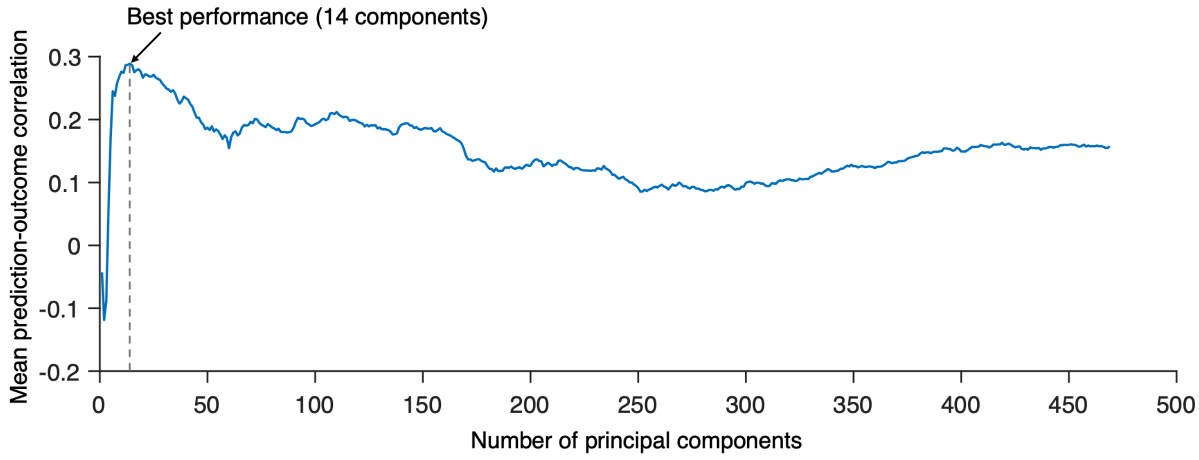


**Figure S6. Predictive performances across different numbers of PCs.**

We tested PCR with a different number of PCs to find the best model to predict the within-individual variation of sustained pain ratings and calculated the mean correlation between actual and predicted pain ratings (10 ratings). Predictive performances were based on leave-one-subject-out cross-validation. The best model used 14 PCs for prediction.

**Table S1. Top 50 stable connections of the classification model**

| Rank | Weights | ROI names | MNI coordinates |
|---|---|---|---|
| *Positive connections* | | | |
| #1 | 0.0222 | LH_SomMot_6 - BG_L_6_6 (dlPu) | (-56,-8,30) - (-28,-6,2) |
| #2 | 0.0200 | LH_SomMot_6 - BG_R_6_6 (dlPu) | (-56,-8,30) - (30,-4,2) |
| #3 | 0.0200 | RH_SomMot_7 - BG_L_6_6 (dlPu) | (58,-4,30) - (-28,-6,2) |
| #4 | 0.0181 | LH_SomMot_6 - BG_L_6_2 (GP) | (-56,-8,30) - (-22,-2,4) |
| #5 | 0.0170 | RH_SomMot_7 - BG_R_6_6 (dlPu) | (58,-4,30) - (30,-4,2) |
| #6 | 0.0162 | LH_SomMot_7 - BG_R_6_6 (dlPu) | (-48,-8,46) - (30,-4,2) |
| #7 | 0.0161 | RH_SomMot_7 - BG_L_6_2 (GP) | (58,-4,30) - (-22,-2,4) |
| #8 | 0.0160 | LH_SomMot_7 - BG_L_6_6 (dlPu) | (-48,-8,46) - (-28,-6,2) |
| #9 | 0.0156 | RH_DorsAttn_Post_9 - RH_SalVentAttn_TempOccPar_2 | (8,-56,62) - (60,-38,16) |
| #10 | 0.0154 | LH_SomMot_6 - BG_R_6_2 (GP) | (-56,-8,30) - (22,-2,4) |
| #11 | 0.0154 | RH_SalVentAttn_PrC_1 - RH_SalVentAttn_FrOperIns_3 | (50,4,40) - (36,24,4) |
| #12 | 0.0154 | LH_Cont_Par_3 - RH_SalVentAttn_Med_3 | (-46,-42,46) - (8,4,66) |
| #13 | 0.0153 | LH_SomMot_2 - BG_R_6_6 (dlPu) | (-52,-24,10) - (30,-4,2) |
| #14 | 0.0161 | LH_SomMot_6 - Cb_Right_VIIb | (-56,-8,30) - (-24,-58,-52) |
| #15 | 0.0160 | RH_SomMot_1 - BG_L_6_6 (dlPu) | (52,-14,6) - (-28,-6,2) |
| #16 | 0.0156 | LH_DorsAttn_FEF_1 - RH_Cont_PFCl_5 | (-32,-4,54) - (30,48,28) |
| #17 | 0.0154 | LH_SomMot_6 - Tha_L_8_2 (mPMtha) | (-56,-8,30) - (-18,-14,4) |
| #18 | 0.0154 | LH_SomMot_6 - Cb_Right_VI | (-56,-8,30) - (24,-58,-26) |
| #19 | 0.0153 | RH_SomMot_1 - BG_R_6_6 (dlPu) | (52,-14,6) - (30,-4,2) |

(Table S1 continues on next page)

| Rank | Weights | ROI names | MNI coordinates |
|---|---|---|---|
| | | | (Continued from previous page) |
| #20 | 0.0139 | RH_SomMot_2 - BG_R_6_6 (dlPu) | (64,-24,8) - (30,-4,2) |
| #21 | 0.0133 | LH_SomMot_2 - BG_R_6_2 (GP) | (-52,-24,10) - (22,-2,4) |
| #22 | 0.0131 | RH_SomMot_2 - BG_R_6_2 (GP) | (64,-24,8) - (22,-2,4) |
| #23 | 0.0130 | RH_SomMot_7 - BG_R_6_2 (GP) | (58,-4,30) - (22,-2,4) |
| #24 | 0.0127 | LH_SomMot_7 - BG_L_6_2 (GP) | (-48,-8,46) - (-22,-2,4) |
| #25 | 0.0127 | LH_SomMot_6 - Cb_Left_VI | (-56,-8,30) - (-22,-58,-24) |
| #26 | 0.0125 | RH_SomMot_10 - BG_R_6_6 (dlPu) | (46,-12,48) - (30,-4,2) |
| #27 | 0.0118 | LH_SomMot_6 - Cb_Left_VIIb | (-56,-8,30) - (-26,-66,-50) |
| #28 | 0.0113 | LH_SomMot_6 - Tha_R_8_8 (lPFtha) | (-56,-8,30) - (12,-16,6) |
| #29 | 0.0112 | LH_SomMot_7 - BG_R_6_2 (GP) | (-48,-8,46) - (22,-2,4) |
| #30 | 0.0111 | LH_Vis_8 - Tha_L_8_2 (mPMtha) | (-48,-70,10) - (-18,-14,4) |
| #31 | 0.0104 | LH_SomMot_13 - LH_Default_PFC_7 | (-26,-38,68) - (-8,58,20) |
| #32 | 0.0104 | LH_Vis_8 - LH_Cont_Cing_1 | (-48,-70,10) - (-4,-28,26) |
| #33 | 0.0103 | RH_SomMot_2 - Tha_L_8_2 (mPMtha) | (64,-24,8) - (-18,-14,4) |
| #34 | 0.0095 | LH_SomMot_13 - LH_Default_PFC_2 | (-26,-38,68) - (-6,36,-10) |
| #35 | 0.0092 | LH_SomMot_7 - Cb_Right_VI | (-48,-8,46) - (24,-58,-26) |
| #36 | 0.0092 | LH_SomMot_13 - RH_Default_PFCdPFCm_1 | (-26,-38,68) - (4,36,-14) |
| #37 | 0.0077 | LH_Cont_Cing_1 - RH_Vis_5 | (-4,-28,26) - (48,-72,-6) |
| #38 | 0.0067 | LH_SomMot_13 - LH_Limbic_OFC_2 | (-26,-38,68) - (-10,36,-20) |

(Table S1 continues on next page)

| Rank | Weights | ROI names | MNI coordinates |
|---|---|---|---|
| | | | (Continued from previous page) |
| *Negative connections* | | | |
| #1 | -0.0235 | RH_SomMot_7 - RH_SomMot_12 | (58,-4,30) - (40,-24,58) |
| #2 | -0.0231 | LH_SomMot_6 - RH_SomMot_12 | (-56,-8,30) - (40,-24,58) |
| #3 | -0.0160 | LH_SomMot_4 - RH_SomMot_12 | (-54,-4,10) - (40,-24,58) |
| #4 | -0.0145 | RH_SomMot_6 - RH_SomMot_14 | (56,-12,14) - (32,-22,64) |
| #5 | -0.0143 | LH_SomMot_6 - LH_Default_Temp_3 | (-56,-8,30) - (-56,-6,-12) |
| #6 | -0.0141 | LH_SomMot_6 - LH_Default_Temp_4 | (-56,-8,30) - (-58,-30,-4) |
| #7 | -0.0135 | LH_SomMot_6 - LH_Default_pCunPCC_1 | (-56,-8,30) - (-12,-56,14) |
| #8 | -0.0124 | LH_SomMot_4 - LH_SomMot_13 | (-54,-4,10) - (-26,-38,68) |
| #9 | -0.0115 | LH_SomMot_4 - RH_SomMot_14 | (-54,-4,10) - (32,-22,64) |
| #10 | -0.0092 | RH_SalVentAttn_Med_1 - Cb_Left_VIIIb | (8,8,42) - (0,-64,-42) |
| #11 | -0.0084 | LH_SalVentAttn_FrOperIns_4 - Cb_Left_VIIIb | (-52,8,10) - (0,-64,-42) |
| #12 | -0.0083 | LH_SalVentAttn_Med_1 - Cb_Left_VIIIb | (-6,10,42) - (0,-64,-42) |

*Note.* Top 50 stable connections based on bootstrap tests with 10,000 iterations (edge-level $P < 1.9 \times 10^{-7}$, FDR $q < 1.4 \times 10^{-4}$).

**Table S2. Top 50 stable connections of the regression model**

| Rank | Weights | ROI names | MNI coordinates |
|---|---|---|---|
| *Positive connections* | | | |
| #1 | 0.000496 | LH_SomMot_12 - RH_Default_pCunPCC_3 | (-32,-22,64) - (6,-58,44) |
| #2 | 0.000460 | LH_SomMot_10 - RH_Default_pCunPCC_3 | (-40,-26,58) - (6,-58,44) |
| *Negative connections* | | | |
| #1 | -0.000768 | RH_Cont_Cing_1 - Cb_Left_Crus_II | (6,-26,30) - (-26,-74,-42) |
| #2 | -0.000756 | RH_Cont_Cing_1 - Cb_Left_Crus_I | (6,-26,30) - (-36,-68,-32) |
| #3 | -0.000743 | Tha_R_8_1 (mPFtha) - Cb_Vermis_VI | (8,-10,6) - (0,-70,-22) |
| #4 | -0.000711 | Tha_R_8_1 (mPFtha) - Cb_Vermis_VIIIa | (8,-10,6) - (26,-58,-54) |
| #5 | -0.000697 | RH_Cont_Cing_1 - Cb_Right_Crus_I | (6,-26,30) - (38,-68,-32) |
| #6 | -0.000695 | RH_Cont_Cing_1 - Cb_Vermis_IX | (6,-26,30) - (6,-54,-48) |
| #7 | -0.000694 | Tha_R_8_1 (mPFtha) - Cb_Left_VIIb | (8,-10,6) - (-26,-66,-50) |
| #8 | -0.000673 | RH_Cont_Cing_1 - Cb_Right_Crus_II | (6,-26,30) - (26,-76,-42) |
| #9 | -0.000662 | Tha_R_8_1 (mPFtha) - Cb_Left_Crus_I | (8,-10,6) - (-36,-68,-32) |
| #10 | -0.000655 | Tha_R_8_1 (mPFtha) - Cb_Vermis_IX | (8,-10,6) - (6,-54,-48) |
| #11 | -0.000655 | LH_Cont_Cing_1 - Cb_Left_Crus_I | (-4,-28,26) - (-36,-68,-32) |
| #12 | -0.000654 | RH_Cont_PFCmp_1 - Cb_Left_Crus_I | (8,30,28) - (-36,-68,-32) |
| #13 | -0.000652 | RH_Cont_PFCmp_1 - Cb_Left_Crus_II | (8,30,28) - (-26,-74,-42) |
| #14 | -0.000650 | Tha_R_8_1 (mPFtha) - Cb_Left_Crus_II | (8,-10,6) - (-26,-74,-42) |
| #15 | -0.000648 | LH_Cont_Cing_1 - Cb_Left_Crus_II | (-4,-28,26) - (-26,-74,-42) |

(Table S2 continues on next page)

| Rank | Weights | ROI names | MNI coordinates |
|---|---|---|---|
| | | | (Continued from previous page) |
| #16 | -0.000647 | Tha_L_8_1 (mPFtha) - Cb_Vermis_VI | (-6,-12,6) - (0,-70,-22) |
| #17 | -0.000643 | LH_Cont_Cing_1 - Cb_Right_Crus_I | (-4,-28,26) - (38,-68,-32) |
| #18 | -0.000642 | Tha_R_8_1 (mPFtha) - Cb_Right_VIIb | (8,-10,6) - (-24,-58,-52) |
| #19 | -0.000626 | BG_R_6_1 (vCa) - Cb_Vermis_VI | (14,14,-2) - (0,-70,-22) |
| #20 | -0.000624 | LH_Cont_Cing_1 - Cb_Right_Crus_II | (-4,-28,26) - (26,-76,-42) |
| #21 | -0.000598 | Tha_R_8_1 (mPFtha) - Cb_Right_VI | (8,-10,6) - (24,-58,-26) |
| #22 | -0.000592 | Tha_R_8_1 (mPFtha) - Cb_Right_Crus_I | (8,-10,6) - (38,-68,-32) |
| #23 | -0.000578 | RH_SalVentAttn_FrOperIns_3 - Cb_Left_VIIb | (36,24,4) - (-26,-66,-50) |
| #24 | -0.000565 | LH_SalVentAttn_Med_1 - Cb_Left_VIIb | (-6,10,42) - (-26,-66,-50) |
| #25 | -0.000549 | Tha_L_8_7 (cTtha) - Cb_Left_Crus_I | (-10,-22,14) - (-36,-68,-32) |
| #26 | -0.000543 | Tha_L_8_7 (cTtha) - Cb_Left_VIIb | (-10,-22,14) - (-26,-66,-50) |
| #27 | -0.000538 | BG_L_6_5 (dCa) - Cb_Vermis_VIIIa | (-14,2,16) - (26,-58,-54) |
| #28 | -0.000532 | LH_SalVentAttn_Med_1 - Cb_Right_VIIb | (-6,10,42) - (-24,-58,-52) |
| #29 | -0.000530 | BG_L_6_5 (dCa) - Cb_Vermis_VI | (-14,2,16) - (0,-70,-22) |
| #30 | -0.000530 | Tha_L_8_7 (cTtha) - Cb_Left_Crus_II | (-10,-22,14) - (-26,-74,-42) |
| #31 | -0.000523 | BG_R_6_5 (dCa) - Cb_Left_Crus_I | (14,6,14) - (-36,-68,-32) |
| #32 | -0.000520 | BG_R_6_5 (dCa) - Cb_Left_VIIb | (14,6,14) - (-26,-66,-50) |
| #33 | -0.000519 | Tha_L_8_1 (mPFtha) - Cb_Left_Crus_I | (-6,-12,6) - (-36,-68,-32) |
| #34 | -0.000515 | BG_R_6_5 (dCa) - Cb_Left_Crus_II | (14,6,14) - (-26,-74,-42) |

(Table S2 continues on next page)

| Rank | Weights | ROI names | MNI coordinates |
|---|---|---|---|
| | | | (Continued from previous page) |
| #35 | -0.000494 | BG_L_6_5 (dCa) - Cb_Left_VIIb | (-14,2,16) - (-26,-66,-50) |
| #36 | -0.000481 | BG_L_6_4 (vmPu) - Cb_Right_VI | (-22,6,-4) - (24,-58,-26) |
| #37 | -0.000477 | Tha_R_8_4 (rTtha) - Cb_Left_Crus_II | (2,-12,6) - (-26,-74,-42) |
| #38 | -0.000473 | Tha_R_8_4 (rTtha) - Cb_Left_Crus_I | (2,-12,6) - (-36,-68,-32) |
| #39 | -0.000470 | BG_L_6_5 (dCa) - Cb_Left_Crus_I | (-14,2,16) - (-36,-68,-32) |
| #40 | -0.000469 | BG_R_6_5 (dCa) - Cb_Right_Crus_I | (14,6,14) - (38,-68,-32) |
| #41 | -0.000454 | BG_R_6_5 (dCa) - Cb_Right_VIIb | (14,6,14) - (-24,-58,-52) |
| #42 | -0.000448 | BG_L_6_5 (dCa) - Cb_Left_Crus_II | (-14,2,16) - (-26,-74,-42) |
| #43 | -0.000444 | RH_Cont_PFCv_1 - Cb_Left_Crus_II | (34,22,-8) - (-26,-74,-42) |
| #44 | -0.000443 | BG_L_6_5 (dCa) - Cb_Right_VIIb | (-14,2,16) - (-24,-58,-52) |
| #45 | -0.000434 | BG_R_6_5 (dCa) - Cb_Right_VI | (14,6,14) - (24,-58,-26) |
| #46 | -0.000421 | BG_L_6_5 (dCa) - Cb_Right_Crus_I | (-14,2,16) - (38,-68,-32) |
| #47 | -0.000405 | BG_L_6_5 (dCa) - Cb_Right_VI | (-14,2,16) - (24,-58,-26) |
| #48 | -0.000393 | BG_L_6_5 (dCa) - Cb_Left_VI | (-14,2,16) - (-22,-58,-24) |

*Note.* Top 50 stable connections based on bootstrap tests with 10,000 iterations (edge-level $P < 6.1 \times 10^{-5}$, FDR $q < 0.043$).
